# Impacts of recreational angling on fish population recovery after a commercial fishing ban

**DOI:** 10.1101/2022.03.07.483248

**Authors:** Justas Dainys, Eglė Jakubavičiūtė, Harry Gorfine, Mindaugas Kirka, Alina Raklevičiūtė, Augustas Morkvėnas, Žilvinas Pūtys, Linas Ložys, Asta Audzijonyte

## Abstract

It is often assumed that recreational fishing has negligible impact on fish stocks compared to commercial fishing. Yet, for inland water bodies in densely populated areas, this is unlikely to be true. In this study we demonstrate remarkably variable stock recovery rates among different fish species with similar life histories in a large productive inland freshwater ecosystem (Kaunas Reservoir, Lithuania), where all commercial fishing has been banned since 2013. We conducted over 900 surveys of recreational anglers during a period of four years (2016 to 2021) to assess recreational fishing catches. These surveys are combined with drone and fishfinder device-based assessment of recreational fishing effort. Fish population recovery rates were assessed using standardised catch per unit effort time series. We show that recreational fishing is having a major impact in retarding the recovery of predatory species, such as pikeperch and perch. In contrast, recovery of roach, rarely caught by anglers, has been remarkably rapid and the species is now dominating the ecosystem. Our study demonstrates that recreational fishing can have strong impacts on some fish species, alter relative species composition and potentially change ecosystem state and dynamics.

## 1. Introduction

Lakes and rivers have been an essential source of fish and other ecosystem services for people all around the world. Yet, these are also among the most vulnerable ecosystems, heavily impacted by fishing, pollution, energy generation and other human activities. Despite freshwater fish populations being among the most threatened in the world (Su *et al*., 2021), the status of fish stocks in inland waters has received far less consideration than marine stocks (FAO, 1999; Hilborn *et al*., 2003; Kura *et al*., 2004; Funge-Smith & Bennett, 2019). This is partly due to limited funding because most commercially valuable and therefore properly researched fisheries occur in marine ecosystems (Walters, 1998; Mangin *et al*., 2018). Another reason is that commercial fishing, which is often easier to monitor, has largely diminished in inland waters (FAO, 2010), while at the same time, the effort from more participants engaging in freshwater recreational fishing has increased rapidly (Cowx, 2002; Coleman *et al*., 2004). The global recreational harvest is poorly documented but may be in the order of 2 million metric tons (FAO, 1999), yet there are few studies which specifically address the influence of recreational fishing on stock recovery after commercial fishing closures.

In developed countries, recreational angling often involves participation rates of 10–30% of the total population (Arlinghaus & Cooke 2009; Arlinghaus *et al*., 2015), constituting considerable social and economic activity and potentially exerting substantial environmental pressure. This is not surprising, considering that about half the people around the globe live within several kilometers of a freshwater body (Kummu *et al*., 2011). Moreover, during recent decades recreational angling has also become more efficient, facilitated by the use of advanced fish finding devices and complex recreational gear, as well as increased mobility of anglers in search of fishing opportunities (Papenfuss *et al*., 2015). Yet, the sociopolitical importance of recreational angling and a lack of accurate estimates to draw attention to its impact (Wadiwel, 2019) means that its sustainability is seldom questioned or seriously addressed (McPhee *et al*., 2002). The prevailing perception among many people is that angling is a relatively ecologically benign activity (Kearney, 2001) which has negligible impact compared to commercial fishing.

Estimation of the total recreational catch and rate of fishing mortality is important both ecologically, and from a regulatory perspective (Pope *et al*., 2017), but it can also be politically contentious (Winstanley, 2019). Management agencies have increasingly relied on size, daily bag or trip limits, and seasonal closures to manage recreational catches (Bartholomew & Bohnsack, 2005). Some studies demonstrated that citizen science can also be used for fish population size structure estimates (Conron *et al*., 2012).

Estimates of recreational fishing mortality often rely on tagging or post-release survival (Vandergroot, 2014, Kristensen *et al*., 2020), but are rarely available for inland ecosystems. In Lake Erie (USA) commercial estimates of Walleye fishing mortality over the course of 18 years ranged around 0.20 to a peak of 0.55, whereas recreational fishing mortality was more stable over time and ranged between 0.05–0.16 (Vandergroot, 2014). Yet, in nearby Lakes St Clair and Huron recreational fishing mortality reached 0.37 in 2005. Other studies also showed that anglers can be highly efficient, with e.g. nearly 20% of pike repeatedly caught by anglers within a 1.5-year period in Lake Mjels in Denmark (Kristensen *et al*., 2020). In general, there are many cases in which recreational catches exceed those taken by the commercial sector (Coleman *et al*., 2004; Cooke & Cowx, 2004; Morales-Nin *et al*., 2005) and where the negative effects of recreational fishing on aquatic systems have been clearly demonstrated (West & Gordon, 1994; McPhee *et al*., 2002; Coleman *et al*., 2004; Cooke & Cowx, 2004).

Kaunas Reservoir is a highly productive and relatively large (65 km^2^) water body, which for decades supported important commercial fisheries and is also one of the most popular recreational fishing destinations in Lithuania. A commercial fishery operated in the reservoir from the 1960s, with catches averaging ca 100 tons per year but reaching almost 230 tons in the early 2000s. From 2005 onwards, commercial catches steadily declined, with stocks of many species collapsing, leading to the closure of all commercial fisheries in 2013. Regular scientific surveys have been conducted in the reservoir since 1992 and some fish species have shown remarkable rates of recovery over the last decade (Ložys *et al*., 2020). Nevertheless, recovery rates appear to vary greatly across species, posing a question about potential impacts of recreational fishing, especially considering that the reservoir has become one of the most popular angling spots in the country. In this study we assess fish population abundance changes using data from scientific surveys, estimate recreational fishing catches and explore whether different rates of species recovery could be explained by mortality imposed by recreational fishing.

## 2. Materials and methods

### 2.1 Research area and commercial catch data

Our study area is Kaunas Reservoir (54.87, 24.14), Lithuania’s largest artificial water body, created in 1959 (Fig. 1). It occupies 65 km^2^, spans 3.3 km at its widest point, and has a maximum depth of 22 meters. The reservoir is a highly productive ecosystem and for decades supported an intensive commercial fishery, with annual catches reaching a peak in year 2004 (total catch of all species was 229 t) (Fig. A.1). Based on scientific surveys the dominant fish species are roach (*Rutillus rutillus*), bream (*Abramis brama*), pikeperch (*Sander lucioperca*), perch (*Perca fluviatilis*) and silver bream (*Blicca bjoerkna*) (Ložys *et al*., 2020), however abundance of some species like Prussian carp (*Carassius gibelio*), carp (*Cyprinus carpio*) and vimba (*Vimba vimba*) might be underestimated due to scientific survey gillnetting methods (see below). Data about commercial catch were collected from monthly catch statistics reported by the commercial fishing bodies and compiled by the Ministry of Environment or similar institutions since 1961. Since 2013 all commercial fishing has been banned.

**Figure 1.**
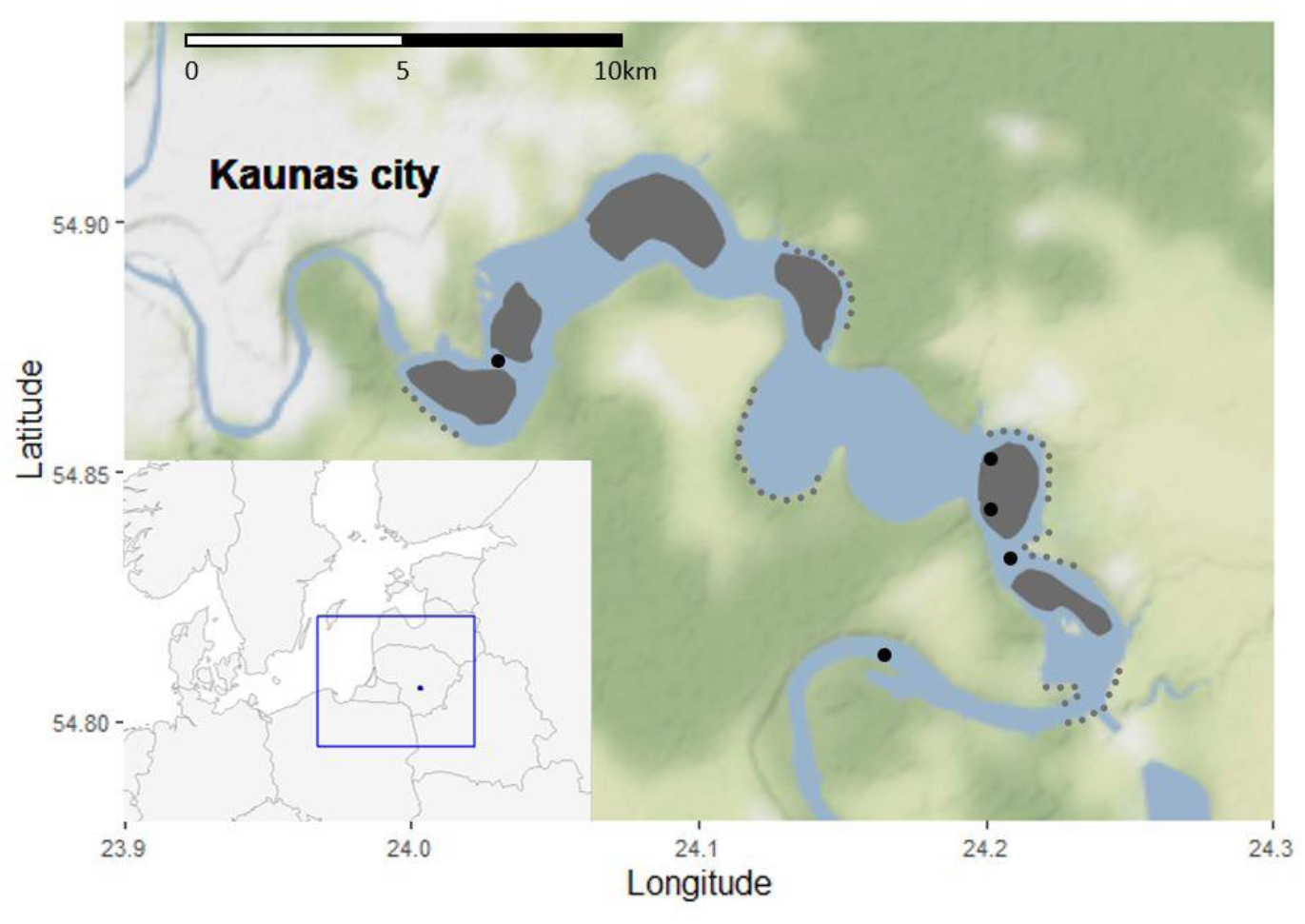
Map of Kaunas Reservoir with grey dotted lines showing locations of angler surveys during the open water fishing season and grey areas – surveys during the ice fishing season. Black dots represent locations of scientific surveys.

### 2.2 Scientific fish abundance data collection

Scientific sampling in Kaunas Reservoir was conducted during 1991–2021 using a range of standard gillnetting approaches. In most cases, two sets of selective gillnet series ranging from 14 to 70 mm (14, 17, 21.5, 25, 30, 33, 38, 45, 50, 60, and 70 mm (knot to knot)) mesh size were used at each sampling station. The nets were typically set between 17:00 and 19:00 h and lifted the next day between 07:00 and 09:00 h, although different soak times have been used during the 30-year sampling period. Net lengths differed among sampling periods and this effect was accommodated in the statistical catch per unit effort (CPUE) standardisation (below). Catches were typically weighed to the nearest gram, separately for each mesh size, net and species. In aggregate, the study included 480 fishing events.

### 2.3 Statistical analyses of scientific monitoring data

Standardised annual CPUE was estimated for five main fish species commonly observed during scientific surveys – roach, bream, silver bream, perch, and pikeperch. Other fish species, such as vimba, Prussian carp or carp are also commonly caught by anglers in Kaunas Reservoir, but their abundances cannot be estimated using gillnetting surveys. This is either because these species mostly stay in shallow vegetated areas which cannot be sampled with gillnets or, in the case of carp, because even the largest mesh of 70 mm is too small. To account for systematic and random variation across sampling events, we standardised CPUE for each of the five fish species using generalised linear models (GLM), as implemented in the R package ‘statmod’ (v. 1.4.33; Giner & Smyth, 2016). For the standardisation, year, season, gear material (nylon, capron), gear length, soak time and mesh size were treated as fixed effects, and residuals of the year parameter were used as standardised CPUE values:

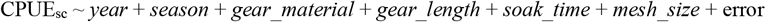

To account for the statistical properties of the scientific survey data, where many fishing events yield zero catches for a given species, we used the Tweedie distribution (Shono, 2008), as implemented in the R package ‘tweedie’ (v. 2.3.3; Dunn, 2017). For each of the species, the analyses were initiated using the full model which included all six potentially explanatory variables, and then the parameter number was reduced based on Akaike’s Information Criterion (AIC; Akaike 1974). The model with lowest AIC score was selected as the best model (Table A. 1), as is commonly used in scientific CPUE standardisation routines (e.g., Forrestal *et al*., 2019). The parameter *year* was treated as an unordered factor, which means that for each *year* value the model estimated separate coefficients. This series of *year* deviations was then extracted as GLM model coefficients and used in further analyses as standardised annual CPUE. For visual purposes and easier comparison across species, these CPUE values were scaled to ensure that the mean estimate in the entire time series is equal to 1, using the R package ‘r4cpue’ (Haddon, 2020). Exact net lengths, mesh sizes or soak time were not available for all scientific survey data. Therefore, in GLM analyses net lengths were assigned into five groups (short to very long), mesh sizes into three groups (small, large, and full), and soak times into three values (short to long) and treated as ordered factors. Our sensitivity analyses showed that these categories were sufficient to capture CPUE trends (Table A.2., Fig. A.2.).

Next, we aimed to assess whether there were significant changes in CPUE associated with the closure of commercial fisheries in 2013. For this we conducted separate CPUE standardisation analyses with the same model parameters included, but in this instance using only data from 2013–2021 and treating *year* as a numeric variable. With this approach we addressed the question of whether there was a linear trend in annual deviations. To assess if there were changes in relative fish abundance across all size classes or only in commercially fished sizes, these analyses were repeated twice. In the first analysis the model included all records from scientific surveys (mesh sizes 14 to 70 mm), whereas in the second analysis only used records for mesh sizes >38 mm, corresponding to commercially used sizes.

### 2.4 On-site recreational fishing surveys

Over the course of 2016, 2017, 2020 and 2021 a total of 910 angler surveys were conducted, with a daily maximum of nearly 150 surveys. Surveys were conducted during all seasons, but the largest number of surveys was completed during the ice fishing season when angler activity was highest (Fig. A.3.). The goal of the angling surveys was to assess recreational catches rather than to estimate recreational effort, which instead was estimated using a different method (see below). We surveyed anglers using slightly modified methods recommended by Malvestuto (1996) and Kaemingk *et al*. (2018). Generally, surveys were distributed throughout the year, with more surveys conducted during weekends and the ice fishing season, when fishing activity was high. During the open water fishing season anglers were interviewed at boat ramps and access points around Kaunas Reservoir, whereas during the ice fishing season anglers were interviewed at their fishing spots on the ice (Fig. 1). Accordingly, in subsequent analyses, fishing trips were categorised as those conducted from the shore, a boat or from ice. Boat angler interviews were mostly complete trips, whereas shoreline interviews comprised both incomplete and complete trips (duration of the fishing trip was included in the model). As recommended by VanDeValk *et al*. (2007) for angling parties containing more than one individual, catch data were collected separately from each angler.

Catch, effort and harvest data were obtained by asking a series of structured questions. The following data were collected during each interview: geographic coordinates of angling location; fishing method (shore, boat, ice); duration of the angling trip; numbers (and weight when possible) and species of fish caught in total; number and species of fish released; numbers, size and species of fish retained; quantity and type of gear used (number of rods per angler); and a few other questions related to gender, age, angling importance and usage of fishfinder devices (not used in analyses presented here) (see Table A.3.). Whenever anglers agreed, the harvested fish were identified, counted, and measured by the interviewers.

### 2.5 Assessment of recreational fishing effort

To estimate total annual recreational catch in Kaunas Reservoir we used two sources of information. First, during 2020 we conducted drone-based angler surveys and calibrated anonymous daily usage of a popular fishfinder device Deeper^®^. Full methodological details, estimates of angler numbers, and uncertainty ranges are available in Dainys *et al*. (under review, provided as a supplement). No recreational effort estimates were available for years prior to 2020. To estimate approximate and relative change in angler effort during earlier years we used data about recreational fishing licence sales, available from the Lithuanian Ministry of Environment. These data were available on an annual basis for 2002–2021 and the number of licences issued were recorded only for Lithuania as a whole and not specific locations, but included annual, monthly and two-day licence categories. In 2002 approx. 35 thousand licenses were sold and by 2014 that number had more than doubled. Since 2014 the number issued has been relatively stable with an average of 74.8 thousand per year (min. 70.7; max. 78.5). To assess the approximate change in recreational effort through time, we assumed that the relative proportion of anglers fishing in Kaunas Reservoir remained stable throughout this period, and that trends in licence sales reflected trends in recreational angling effort.

### 2.6 Statistical analyses for recreational catches

As with the scientific survey CPUE, we used GLMs to estimate recreational CPUE. Here the full CPUE model included year, season, gear type (4 categories: float fishing, feeder (bottom) fishing, spin fishing, ice fishing), number of gears, duration of the fishing trip and method (3 categories: shore, boat, ice).

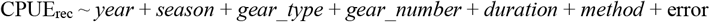

Separate CPUE models were built for each species and for all species combined (total catch per trip). Analyses were done using the Tweedie distribution to account for zero catches. For each species the full model was progressively reduced based on AIC values.

In these analyses we included all reported catch, regardless of whether the fish was retained or released. This was for two reasons. First, it is not known how many of the released fish survive, with estimated release mortality rates ranging from almost zero to nearly 90 % (Muoneke & Childress, 1994; Bartholomew & Bohnsack, 2005). Second, because angler surveys were voluntary and conducted by scientific personnel and not fisheries enforcement officers, some anglers were reluctant to show their catches and responded that all fish were released. Consequently, the retained fish that we measured were likely to be underestimates of the actual retained catches. Future studies are needed to assess the correspondence between voluntarily reported survey catches and those observed by officials with enforcement power; this number is likely to differ across countries and socio-economic groups. To account for both the release mortality and under-reporting, we therefore assumed that all catch was retained, although based on our surveys about 50% of fish caught (by biomass) were released (see Discussion for possible implications of not accounting for released fish).

To estimate total recreational catch, we needed to combine estimates of recreational effort and catch per effort. Here we used estimated daily angler numbers in Kaunas Reservoir, as described in Dainys *et al*. (under review). However, for these daily numbers we did not have information about gear types, numbers, fishing duration or fishing method. Therefore, we developed a set of simplified statistical models to predict recreational CPUE using season as the only predictive variable. For nearly all species, season was one of the most significant factors in the full models (see Results), notwithstanding that the season-only models explained considerably less variation than the full models (see Results). Nevertheless, parameter estimates for full and season-only models were generally similar (Table A.4.), indicating that season-only models provided a reasonable general estimate of recreational catch, given the limited available data.

The R code and full details of the statistical analyses and datasets of recreational data are available as a supplement to this manuscript and through https://github.com/astaaudzi/recreationalCatch. Scientific CPUE standardisation used similar methods and code to the recreational catch CPUE standardisation details.

## 3. Results

### 3.1 Scientific catch-per-unit-effort trends show slow recovery rates of piscivorous species

The standardised scientific CPUE values for bream, silver bream, roach and pikeperch were higher during the 1990s and then decreased in the 2000s (Fig. 2), whereas for perch the values remained relatively stable throughout the 1990s to the 2000s. The full scientific monitoring dataset showed that for roach and silver bream, standardised CPUE has nearly tripled over the last 8 years since the closure of commercial fishery in 2013 (Fig. 2), although there appears to be a decline in roach CPUE during the last few years. In contrast, for bream, perch and pikeperch no such clear trends were evident.

**Figure 2.**
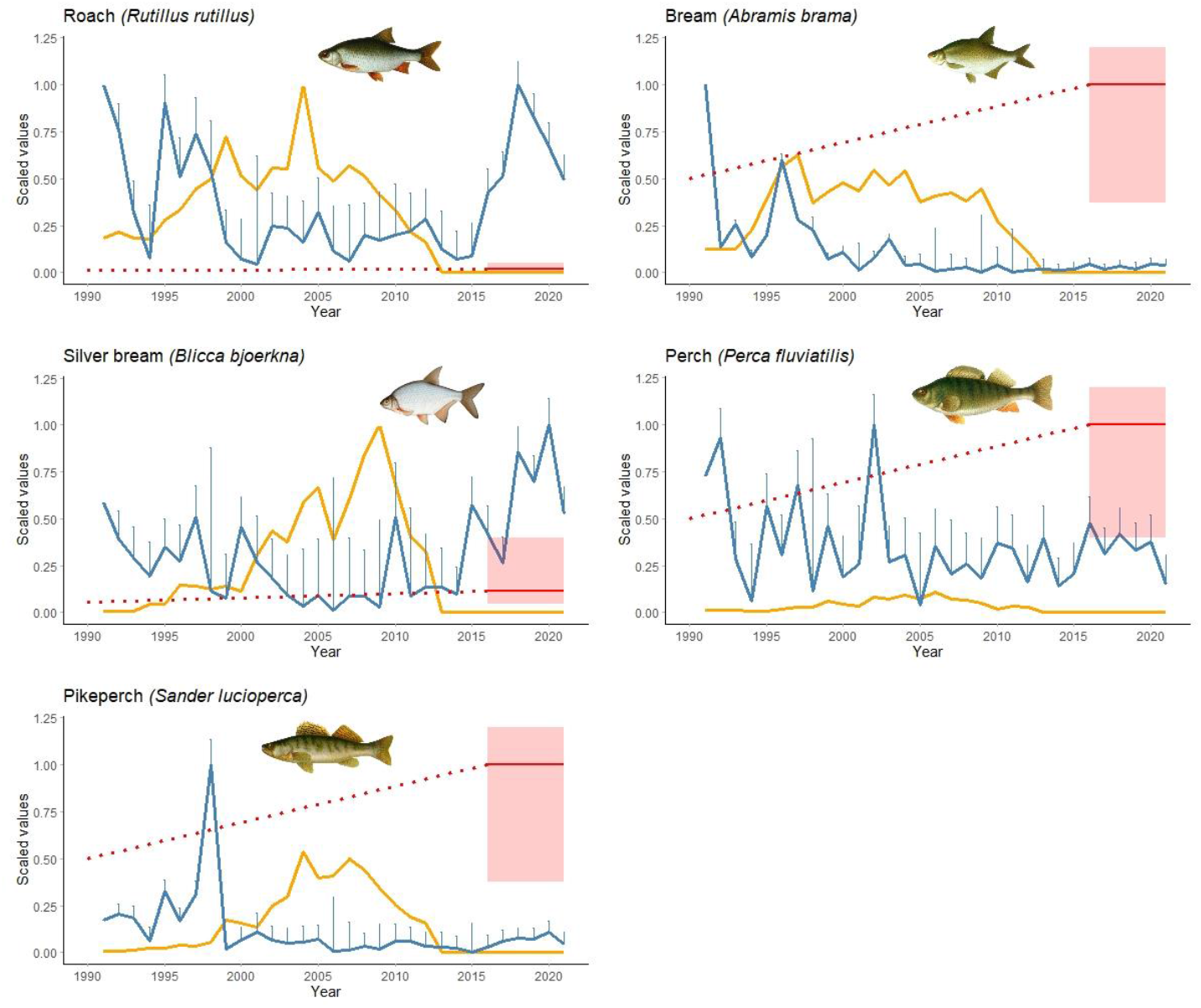
Relative changes in scientific catch-per-unit-effort (CPUE) and commercial and recreational catches for the five main fish species in Kaunas Reservoir. Scientific CPUE show standardized annual values and confidence ranges. Recreational catches were estimated for the last 5 years and show mean estimates and 80% uncertainty ranges. For years since 2016 we provide just an indicative trendline, assuming that recreational effort and therefore catches was about half as intense in 1990 and has been steadily growing since then. The y axis shows relative values, where scientific CPUE was scaled separately, but commercial and recreational catches were scaled together (using mean of estimated recreational catches), so they can be compared. Absolute commercial fishing values are shown in Fig. A.1, estimated for recreational fishing in Table 2.

To assess the magnitude and significance of a linear CPUE trend since 2013, we checked whether there was a significant slope of standardised yearly values in survey data from 2013–2021, where year was treated as a numeric variable (Fig. A.4.). In these analyses roach and silver bream both showed steeply positive (slope of ca 0.2, which means about 20% increase in CPUE per year), highly significant (p < 1e-4) trends. These trends were evident both in the analyses across all fish sizes and those restricted to mesh sizes > 38 mm. Bream also showed a positive, although weaker (ca 10%) slope (p = 0.002 for all sizes and 0.017 for larger sizes). Similarly, a 10% slope was observed in pikeperch when all sizes were included (p = 0.007), but there was no significant trend when only large fish were analysed (p = 0.27). In contrast, there was a negative CPUE trend for perch, suggesting about 6 −7% decrease in scientific CPUE per year, although this trend was only marginally significant for large sizes (p = 0.056).

### 3.2 Recreational angling mostly targets piscivorous fish and some cyprinids

Over the course of 2016, 2017, 2020 and 2021 a total of 910 angler surveys were conducted. In these surveys 53.2 % (N=484) of interviewed anglers were fishing during the open-water season, and from these 484 anglers about one quarter (25.8 %, N=125) were fishing from boats and the rest (74.2 %, N=359) fished from the shore. The remaining 46.8 % of anglers (N=426) were interviewed during the ice fishing season. In total, 14 different fish species were recorded during the surveys: perch (*Perca fluviatilis*), Prussian carp (*Carassius gibelio*), common carp (*Cyprinus carpio*), common bream (*Abramis brama*), common roach (*Rutilus rutilus*), northern pike (*Esox lucius*), silver bream (*Blicca bjoerkna*), ruffe (*Gymnocephalus cernuus*), common rudd (*Scardinius erythrophthalmus*), asp (*Leuciscus aspius*), pikeperch (*Sander lucioperca*), Wels catfish (*Silurus glanis*), chub (*Leuciscus cephalus*) and vimba (*Vimba vimba*). Perch and pikeperch were the main species taken by anglers fishing from boats, comprising almost 79 % of the total catch by numbers. Prussian carp and bream comprised more than half (56.5 %) of the total catch caught by anglers fishing from the shore. Perch was the main species caught by anglers fishing on ice, comprising 57.9 % of the total catch by numbers

In GLMs applied to recreational catch, we found that the best model for the total catch included all parameters (Table A.4.). This model indicated that total recreational CPUE was highest during summer, when fishing from a boat and targeting bottom feeding (demersal) species (Table A.4., Fig. A.5.). The highest catches per trip were recorded in 2020, but there was no obvious trend of increasing catch per trip from 2016 to 2021. Both fishing duration and number of gears used also had a positive and significant effect on the total catch. Total standardised recreational catch per trip was highest in spring and summer (Table 1), reaching approximately 1.0–2.5 kg per fishing trip, and lowest in winter, with about 300 g per trip.

**Table. 1.** Estimated recreational catches (grams per fishing trip) and their uncertainty, for selected species and season combinations, using data from total number of angling trips and models that included only a random intercept and season (for roach season was non-significant).

| Parameter | Mean catch | 2.5% CI | 97.5% CI | *t* | *p* |
| --- | --- | --- | --- | --- | --- |
| **Total catch** |  |  |  |  |  |
| autumn | 991.32 | 752.25 | 1306.37 | 49.06 | 0.00 |
| spring | 1396.70 | 985.08 | 1980.31 | 1.51 | 0.13 |
| summer | 1975.64 | 1551.60 | 2515.58 | 3.69 | 0.00 |
| winter | 299.76 | 211.98 | 423.89 | -5.30 | 0.00 |
| **Roach (*Rutilus rutilus*)** |  |  |  |  |  |
| total average (intercept) | 31.13 | 18.22 | 49.75 | 13.46 | 0.00 |
| **Pikeperch (*Sander lucioperca*)** |  |  |  |  |  |
| autumn | 20.48 | 5.39 | 77.73 | 4.44 | 0.00 |
| spring | 238.11 | 123.95 | 457.43 | 3.24 | 0.00 |
| summer | 302.84 | 188.53 | 486.46 | 3.73 | 0.00 |
| winter | 0.47 | 0.00 | 99.85 | -1.34 | 0.18 |
| **Perch (*Perca fluviatilis*)** |  |  |  |  |  |
| autumn | 140.87 | 88.28 | 224.80 | 20.78 | 0.00 |
| spring | 24.90 | 5.53 | 112.12 | -2.16 | 0.00 |
| summer | 75.37 | 35.98 | 157.87 | -1.40 | 0.00 |
| winter | 170.61 | 124.60 | 233.61 | 0.67 | 0.18 |
| **Bream (*Abramis brama*)** |  |  |  |  |  |
| autumn | 322.90 | 207.51 | 502.45 | 25.64 | 0.00 |
| spring | 430.57 | 242.66 | 763.98 | 0.78 | 0.44 |
| summer | 230.56 | 124.73 | 426.20 | -0.87 | 0.38 |
| winter | 50.22 | 23.75 | 106.18 | -4.20 | 0.00 |
| **Silver bream (*Blica bjoerkna*)** |  |  |  |  |  |
| autumn | 41.37 | 22.43 | 76.31 | 11.93 | 0.00 |
| spring | 3.77 | 0.27 | 53.67 | -1.72 | 0.08 |
| summer | 98.93 | 60.37 | 162.10 | 2.18 | 0.03 |
| winter | 3.17 | 0.76 | 13.20 | -3.25 | 0.00 |
| **Prussian carp (*Carassius gibelio*)** |  |  |  |  |  |
| autumn | 110.61 | 62.77 | 194.90 | 16.30 | 0.00 |
| spring | 643.87 | 440.97 | 940.13 | 5.07 | 0.00 |
| summer | 500.49 | 355.19 | 705.24 | 4.47 | 0.00 |
| winter | 0.00 | 0.00 | Inf | -0.03 | 0.98 |
| **Carp (*Cyprinus carpio*)** |  |  |  |  |  |
| autumn | 220.78 | 112.12 | 434.75 | 15.63 | 0.00 |
| spring | 28.30 | 2.27 | 352.14 | -1.54 | 0.12 |
| summer | 370.37 | 195.08 | 703.16 | 1.09 | 0.28 |
| winter | 0.00 | 0.00 | Inf | -0.02 | 0.99 |
| **Catfish (*Silurus glanis*)** |  |  |  |  |  |
| autumn | 0.00 | 0.00 | Inf | -0.01 | 0.99 |
| spring | 0.00 | 0.00 | Inf | 0.00 | 1.00 |
| summer | 370.37 | 201.49 | 680.79 | 0.01 | 0.99 |
| winter | 0.00 | 0.00 | Inf | 0.00 | 1.00 |

When recreational CPUE was modelled separately for different species, parameter and catch estimates were more uncertain. Nevertheless, in most cases season had a significant effect, with the highest catches of Prussian carp, pikeperch, silver bream, or bream observed in the summer or spring, and boat-based or spin fishing yielding higher catches of predatory species, such as pikeperch and perch (Table A.4., Fig. A.5). For species that were less common in recreational catches, such as roach, pike, asp, vimba and some others, most parameter estimates were highly uncertain. For example, for roach the best recreational CPUE model included only the type of fishing gear, where ice fishing had higher catches than other methods (in this instance although the ice fishing method coincides exclusively with the winter season, catches in other seasons were similar to those in winter and therefore season was not selected as an important variable). At a species level, the highest catches per fishing trip in spring and summer were for Prussian carp, common carp, bream and pikeperch (Table 1). In autumn, the highest recreational CPUE was for bream, carp and perch. In winter, perch dominated catches and uncertainty of parameter estimates for other species was very high due to the limited number of observations.

To assess total catch per year, we combined the estimated annual number of angling trips with the catch per trip. For these extrapolations we used season-only recreational catch CPUE models, because estimates of total angling trips were available daily, but without information about gear types, trip duration or other variables that might affect CPUE. To assess whether season-only CPUE models gave similar magnitude and direction of coefficient estimates, we compared best and season-only CPUE models. Generally, the season coefficient estimates were similar, although these were more uncertain when simpler models were used (Fig. A.5.). This was especially true for catch estimates during winter (for pikeperch, Prussian carp, carp, asp). Season had no significant effect on the catch of roach in both full and season-only models, hence for this species catch extrapolation was undertaken using the intercept only model (Table A.4). Obviously, season-only models explained considerably less variance (5–35%; except for roach, Table A.4), compared to the best model (21–69%), therefore predictions of total catch had high uncertainty ranges (see below).

### 3.3 Estimated recreational catches for some fish species are much larger than former commercial catches

In our earlier study (Dainys *et al*., under review) we estimated the total number of angling trips in Kaunas Reservoir for each season between March 2020 and March 2021. Using Bayesian analyses we showed that during spring, summer and autumn there were (in thousands, K) about 40K (26K–64K), 31K (20K – 49K) and 21K (13K–35K) angling trips, where the ranges in parentheses are the 80% posterior Bayesian probability limits. For the ice-fishing season the estimated that the number of trips was 17K (13K–22K). Combining these estimates with the estimated catch per angling trip and propagating the full error in estimates of numbers of anglers and recreational CPUE estimates, shows that during one year anglers caught about 143 tons (70–305 t) of fish in total (Table 2), with the highest catches observed in summer and spring (61 t and 56 t respectively).

**Table 2.** Estimated annual recreational catches, in tons, for the main fish species in Kaunas Reservoir. For species marked with * the total catches were estimated excluding seasons for which an infinite recreational catch was predicted (see Table A.5.).

| Species | mean | min | max |
| --- | --- | --- | --- |
| Prussian carp | 43.63 | 19.62 | 101.44 |
| Bream | 31.98 | 11.94 | 89.52 |
| Pikeperch* | 19.36 | 7.29 | 54.79 |
| Carp | 17.24 | 5.53 | 72.06 |
| Catfish | 11.49 | 4.12 | 33.37 |
| Perch | 9.09 | 3.61 | 27.9 |
| Silver bream | 4.14 | 1.55 | 14.32 |
| Roach | 3.38 | 1.32 | 8.45 |
| Vimba | 1.29 | 0.36 | 13.45 |
| Asp | 1.07 | 0.32 | 4.08 |
| Pike* | 1.07 | 0.22 | 6.06 |
| **Total catch** | **142.91** | **70.08** | **304.65** |

The annual catch estimates for the five fish species that were important commercially and had scientific CPUE trend estimates were as follows: for pikeperch they were around 19 ton, but with very high uncertainty ranges (7–55 t), for perch at ca 9 t (4–28 t), for bream at ca 32 t (12-90 t), for silver bream at ca 4 t (2–14 t) and only ca 3 t for roach (1–8 t). For roach and silver bream these recreational catches were negligible compared with past commercial catches, which in peak years used to reach >100 t for roach and >30 t for silver bream (Fig. A.1.). For the other species, however, the picture was completely different. For example, for perch, recreational catches were 10–20 times higher than the former commercial take (at least 4 times higher if we apply the minimum uncertainty range of the recreational catch and the peak of the commercial catch). Similarly, for pikeperch the recreational take was about 5 times higher than former commercial catches and for bream about 3 times higher (Fig. 2). Non-commercial fish species like Prussian and common carp also comprised a large proportion of the annual recreational catch, at 44 and 17 t respectively. For these species, commercial annual average catches were 4 and 0.5 t respectively. Although highly uncertain, around 11 t of catfish were caught per year by anglers, whereas annual recreational catches of other species, like asp, pike or vimba, were about 1 t each (Table 2).

## 4. Discussion

Commercial fishing in inland and coastal ecosystems has been in decline (Cowx, 2015). This is generally good news for freshwater ecosystems, given that freshwater fish are among the most vulnerable and threatened taxa in the world (Zhang *et al*., 2020). In line with this general trend, about 10 years ago all commercial fishing ceased due to a regulatory closure in Kaunas Reservoir in Lithuania. It was expected that fish population biomass and abundance would increase rapidly after the closure, but this was only observed in some species, such as roach (Ložys *et al*., 2020). Population growth rates differed greatly among species, but the reasons for this difference remained unclear. This study presents detailed analyses of recreational fishing catches in Kaunas Reservoir and shows that angling, through direct and indirect effects, is likely to be the main reason for variability in responses of fish species to the commercial fishing closure. Although our estimates of recreational anglers’ catches have high uncertainty, they reveal that fish species exhibiting slow or even negative scientific CPUE trends, such as perch or pikeperch, have been subject to high recreational exploitation. This contrasts with the low recreational catch pressure on rapidly recovering species, such as roach or silver bream. Other species, such as Prussian carp or carp may also be adversely affected by angling, but scientific CPUE for these species was imprecise, precluding the detection of trends in abundance. Moreover, both species are non-native to Lithuania and therefore are of less ecological concern.

Although Kaunas Reservoir has always been recognised as a popular angling destination in Lithuania, like many waterbodies in Europe there have been only vague estimates of total recreational catches prior to our study. This has led to a general assumption that recreational angling has a relatively minor impact compared with the commercial sector, and that recreational fishing mortality lies within the range of fluctuations in natural mortality (e.g. below 10% per year). Despite this perception, there is growing recognition that recreational fishing can be a significant contributor to global declines in fish populations, particularly in inland waters (Post *et al*., 2002; Cooke & Cowx, 2004; 2006; Coleman *et al*., 2004). For example, Zeller *et al*. (2008) found that during 1950–2005 non-commercial catches from demersal fish stocks in coastal marine areas in Hawaii were more than double the reported commercial catches. Recreational angling can reduce target fish stock abundance in both marine (e.g., Bennett, 1991) and freshwater (e.g., Olson & Cunningham, 1989; Beard & Essington, 2000; Post *et al*., 2002) systems. Anglers may have a particularly strong influence on depressing the size structure of target fish populations as they exhibit greater size selectivity than commercial fishers, for example through choice of lure and hook size (Beard & Kampa, 1999). For freshwater ecosystems located near densely populated areas, recreational fishing mortality can be extreme, reaching annual exploitation rates up to 80 per cent of the average standing stock biomass (for review see Lewin *et al*., 2006). Post *et al*. (2002) shows that four main Canadian species targeted by anglers (walleye, rainbow trout, lake trout, and pike) became heavily depleted due to recreational fishing, where depensatory mechanisms in population recruitment dynamics possibly led to the collapse of these stocks.

In this study we did not estimate total population biomasses or mortality rates for different fish species in Kaunas Reservoir, as this will require more detailed modelling. Yet it is clear that compared to commercial catches, recreational fishing has an especially strong impact on bream and predatory species, with potentially adverse ecological implications for the entire ecosystem. There might be several reasons why perch and pikeperch recreational catches are considerably higher than past commercial catches. It seems unlikely that perch or pikeperch were not targeted by commercial fishers, because both species are prized and have high resale value. Instead, it seems more likely that angling imposed consistently high mortality rates and as a result, the commercial harvest prior to closure was dominated by other species that were either difficult for recreational anglers to catch (roach) or were not targeted (silver bream). Such a scenario would suggest that anglers were very efficient at targeting perch, pikeperch and bream. This is hardly surprising. Our assessment of recreational effort in Kaunas Reservoir indicates that during 2020 an average of about 300 recreational fishing trips were conducted daily (Dainys *et al*., under review). When combined with angler flexibility to specifically target suitable habitats, technically advanced fishing gear, and a determination to explore a wide range of places, anglers’ catching efficiency is likely to be very high. Yet, some species may still be difficult to catch. Notably, in our interviews many anglers commented that roach would be a desirable catch, yet recreational catches for this species were very low. It is possible that roach are not attracted to lures, due to ample quantities of *Dreissena* mussels, an mollusc which appears to be its main food source, constituting up to 80 % of the roach diet (Yount, 1991). The situation is different for silver bream, another species that has showed rapid recovery since 2013. This species is considered of low value by both recreational and commercial fishers but was a substantial component of bycatch with up to 30 tons per year landed (compared to only about 1 t of recreational catch).

Selective removal of predatory fishes by recreational anglers is a well-recognised issue, which is likely having adverse ecological consequences in Kaunas Reservoir and possibly other Lithuanian and central European lakes and reservoirs. Pikeperch and large perch play important regulatory roles, and their low abundance can lead to trophic cascades where the water body they inhabit becomes dominated by roach and phytoplankton. If biomass and average size of predatory species are severely impacted by anglers, their populations can become less able to exert predatory control on cyprinid abundance. Larger cyprinid species may even switch to feed on perch (Vejřík *et al*., 2016), thereby leading to positive feedback loops in the ecosystem which further exacerbate depletion of perch. Roach, which is extremely abundant in Kaunas Reservoir, is known to be a generalist species (Persson, 1987) that together with other abundant cyprinids like bream and white bream can consume eggs and fry of other species. Direct competition for food and space is also possible, especially at a young age.

Although this study provides the first comprehensive assessment of recreational catches in Lithuania, like all assessments of recreational effort and catch, it has large uncertainty ranges and a range of limitations which warrant some caveats. First, detailed recreational effort data were available only for one year and therefore had to be extrapolated on the basis of recreational fishing licence sales. Such extrapolation is commonly used to estimate recreational effort, but is imprecise. It is possible that during 2020, in association with the outbreak of the COVID-19 pandemic, anglers conducted more trips per annual licence, compared with previous years. This would produce overestimates in our recreational catch estimates for the period 2016–2019. Second, as mentioned in the Methods, we considered that all fish recorded in angler surveys were retained, even though according to angler reports about half of the catch was released. Our choice to ignore the released fish aimed to account for post-release mortality and the likely underestimation of disclosed catches in our voluntary surveys. Nevertheless, even if we consider that catch and release rates were reported accurately and all released fish survived, our recreational catch estimates by biomass would be about 30–50% lower (because most released fish were small, contributing less to the total biomass caught). This would still suggest that recreational catches of perch, pikeperch and bream were far higher than the commercial catches of the past and are likely to be having a major impact on population abundances. Third, for some species our statistical models of recreational CPUE had very high upper confidence ranges, indicating high uncertainty about the maximum possible recreational catch. Yet, the estimated total catch per trip (1.5–2.5kg per trip during the open water season) was compatible with online surveys we conducted earlier and is less than half of the total allowable catch per trip (currently at 5 kg). Access-point creel surveys are generally assumed to provide relatively accurate estimates of angler harvest (Barnes *et al*., 2014). Jones *et al*. (1995) recommended a minimum of 100 interviews for a reliable creel survey, particularly when angler catch rates are highly variable. In this study, we interviewed 910 anglers, which provides fairly strong spatial and temporal coverage. Yet, all of our interviews were done by scientific staff, with voluntary participation (i.e., anglers could refuse to respond to questions). It is normal that harvest data obtained during creel surveys is based on angler reports and is accepted without validation (Lockwood *et al*., 1999; Mallison & Cichra, 2004). Yet, in several lakes in Florida (USA) a comparison between reported and counted harvest revealed differences of 7–22% for anglers targeting sunfish or black crappie, but smaller biases when anglers were targeting sunshine bass, palmetto bass and largemouth bass (Mallison & Cichra, 2004). These biases are likely to differ across countries, recreational fishing rules and socio-economic groups. For example, in the Florida example above, accuracy among species was inversely related to bag limit size (Mallison & Cichra, 2004). In future it would important to assess discrepancies between catches reported based on voluntary angler surveys, such as ours, and those conducted by fisheries regulatory bodies.

Recreational activities such as angling are important for human well-being and can provide substantial contribution to local economies (FAO, 2017; Venohr *et al*., 2018). In many countries, including Lithuania, they also are likely to provide some proportion of the local protein supply. What are the possible management solutions for sustainable use of inland ecosystems? It is unrealistic to expect that low population densities of predatory fishes will automatically reduce angling pressure. Lewin *et al* (2006) point out that perceptions about self-regulation of anglers through reducing fishing pressure in response to declining abundance are false. Instead, awareness campaigns and more targeted management measures may be needed. One of the main challenges in recreational fishing management, is that fishing effort is usually unknown and not regulated. Fishing licenses are not limited, they are cheap and, at least in Lithuania, annual licenses allow an unlimited number of fishing trips. This means that even with regulations on daily bag limits, total catch can be exceptionally high. More successful measures for facilitating resource sustainability and angler satisfaction, are likely to be achieved through size and slot limits, especially if they also protect large and highly fecund individuals (Ahrens *et al*., 2020). Limiting or prohibiting recreational fishing access in protected areas is often contentious and conflict-prone, because peoples’ freedom of choice is affected (Stoll-Kleemann, 2010). However, recreational impacts can be managed using tools that do not prohibit access to a site entirely, such as spatial and temporal zoning (Abell *et al*., 2007; Manning, 2010). In many countries anglers now also engage in catch-and-release fishing, especially for more vulnerable predatory species. This can be an effective measure, although there can be substantial post-release mortality (Muoneke & Childress, 1994) as well as reduced growth and fitness among survivors (Cooke *et al*., 2002). Either way, to understand and assess recreational fishing impacts, society and recreational fisheries managers need reasonably accurate and cost-effective methods to monitor angler effort and catch, as have been initiated during this study and will continue to evolve.

## Supplementary tables

**Supplementary Table A.1**. Categories of net length, mesh sizes and soak time used to standardise scientific catch per unit effort. Because only approximate values were available for some fishing trips, we used 5 net length, 3 mesh size and 3 soak time categories. These categories were sufficient to capture the trend in CPUE, assessed using a smaller number of surveys where exact numeric values of these parameters were available (Fig. A.5)

Gear_lengthcategories:

- “very short” <=20 m,
- “short” – 20-60 m,
- “medium” – 61-120 m,
- “long” – 121-300 m,
- “very long” >300 m.

Mesh_size_ categories:

- “small” <=38 mm,
- “big” >38 mm,
- “full” - when catches were reported for the full range of mesh sizes only (e.g. 17-70 mm).

Soak_time categories:

- “short”, <=5 h,
- “medium” 6-12 h, and
- “long” =>12 h.

**Supplementary Table A.2.**
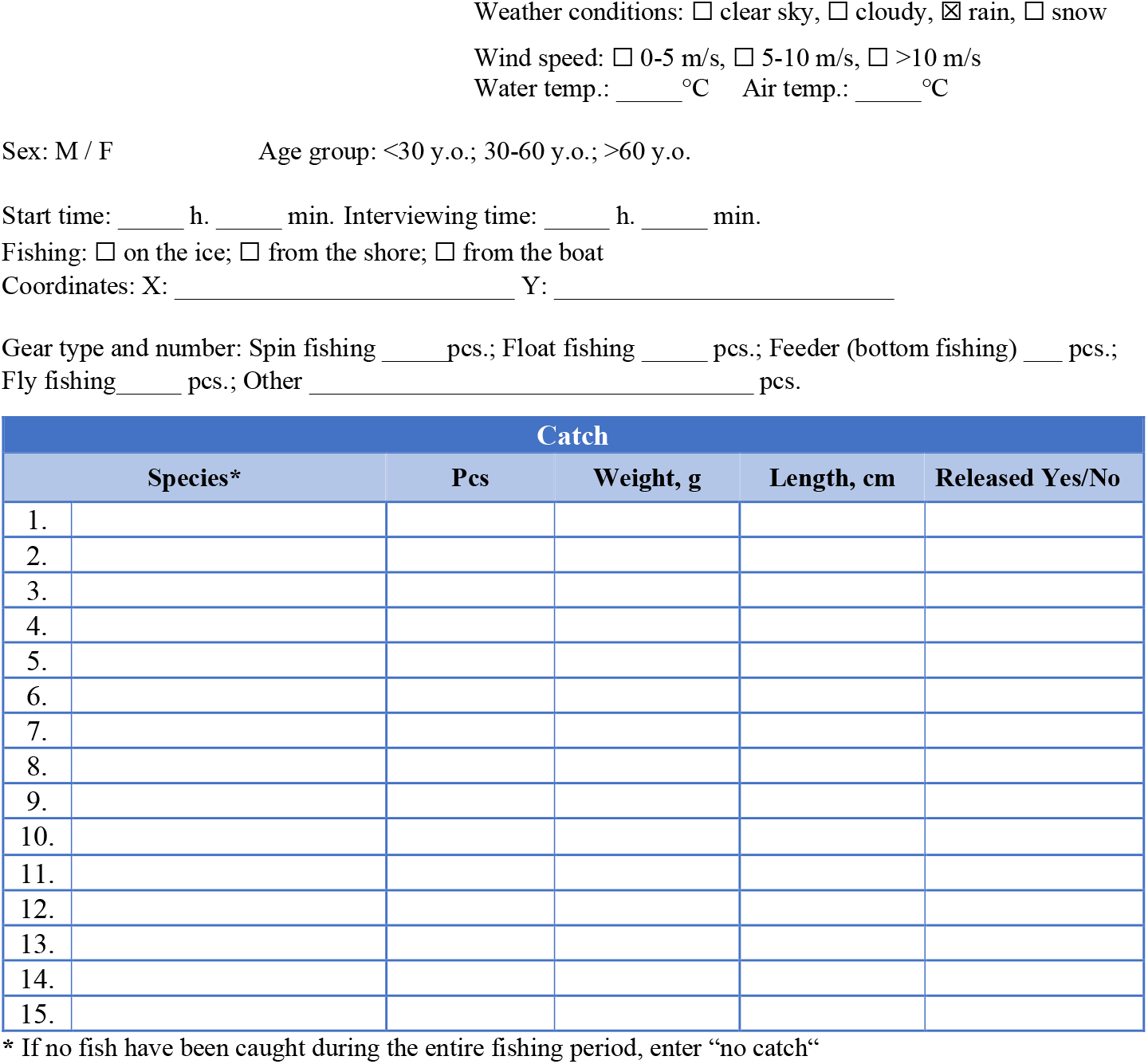
Questionary that was used for onsite angler surveys done by the scientific personnel.

**How important fishing is to you?** □ not very important; □ important; □ very important

**How many times a year do you fish?** □ 1-5; □ 6-10; □ 10-20; □ 20-50; □ >50

**What is the average duration of your fishing trip?** □ <3 h.; □ 3-6 h.; □ 6-9 h.; □ >12 h.

**Do you use fishfinder?** □ Yes; □ No

**If yes, how often?** □ Always; □ 90-50%; □ 50-20%; □ <20% of all fishing trips.

**If yes, how long do you use fishfinder during fishing trip?**

□ only in the beginning (<20% of all fishing time); □ 20-50%, □ 50-100% of all fishing time.

**Supplementary Table A.3.**
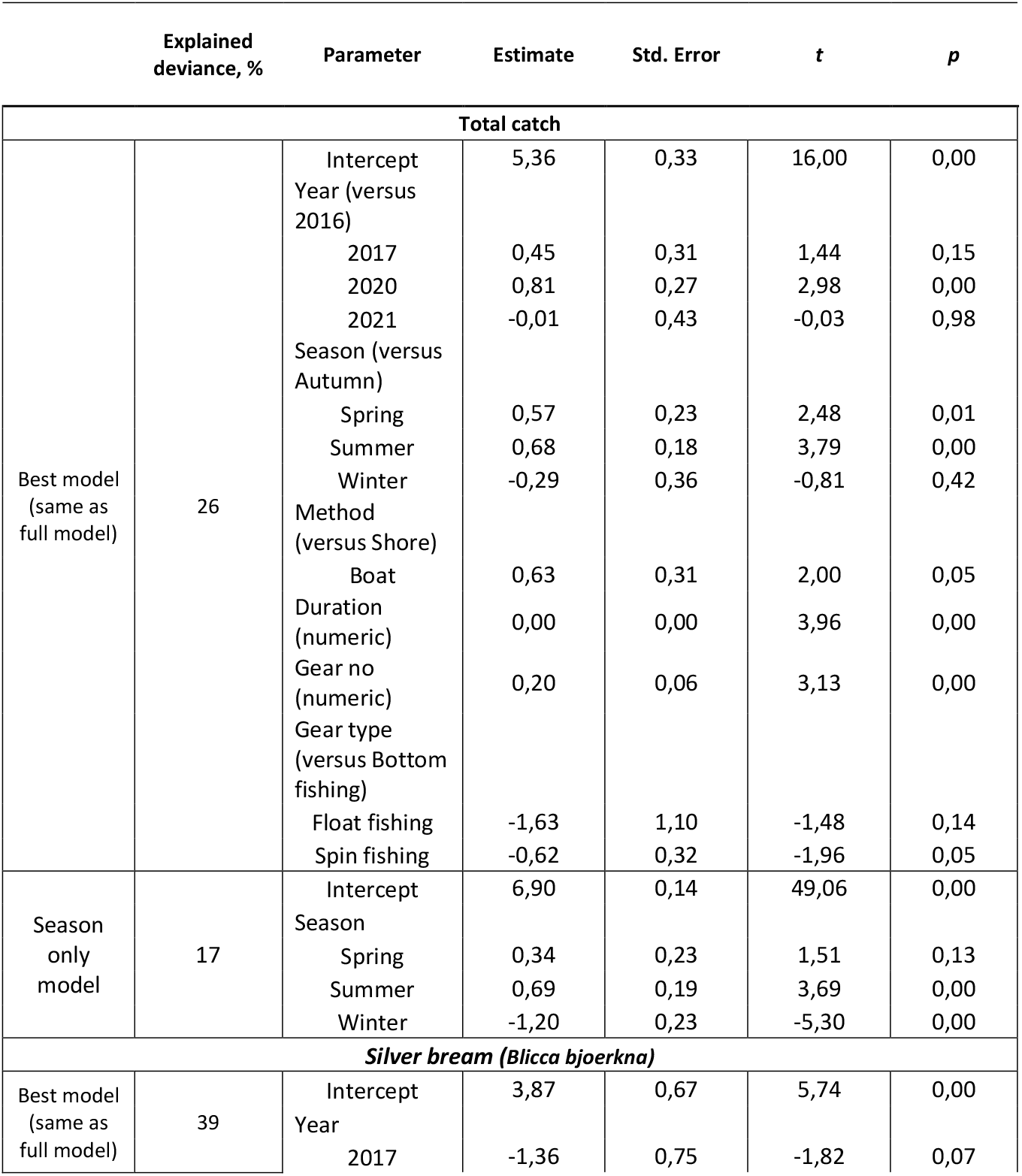
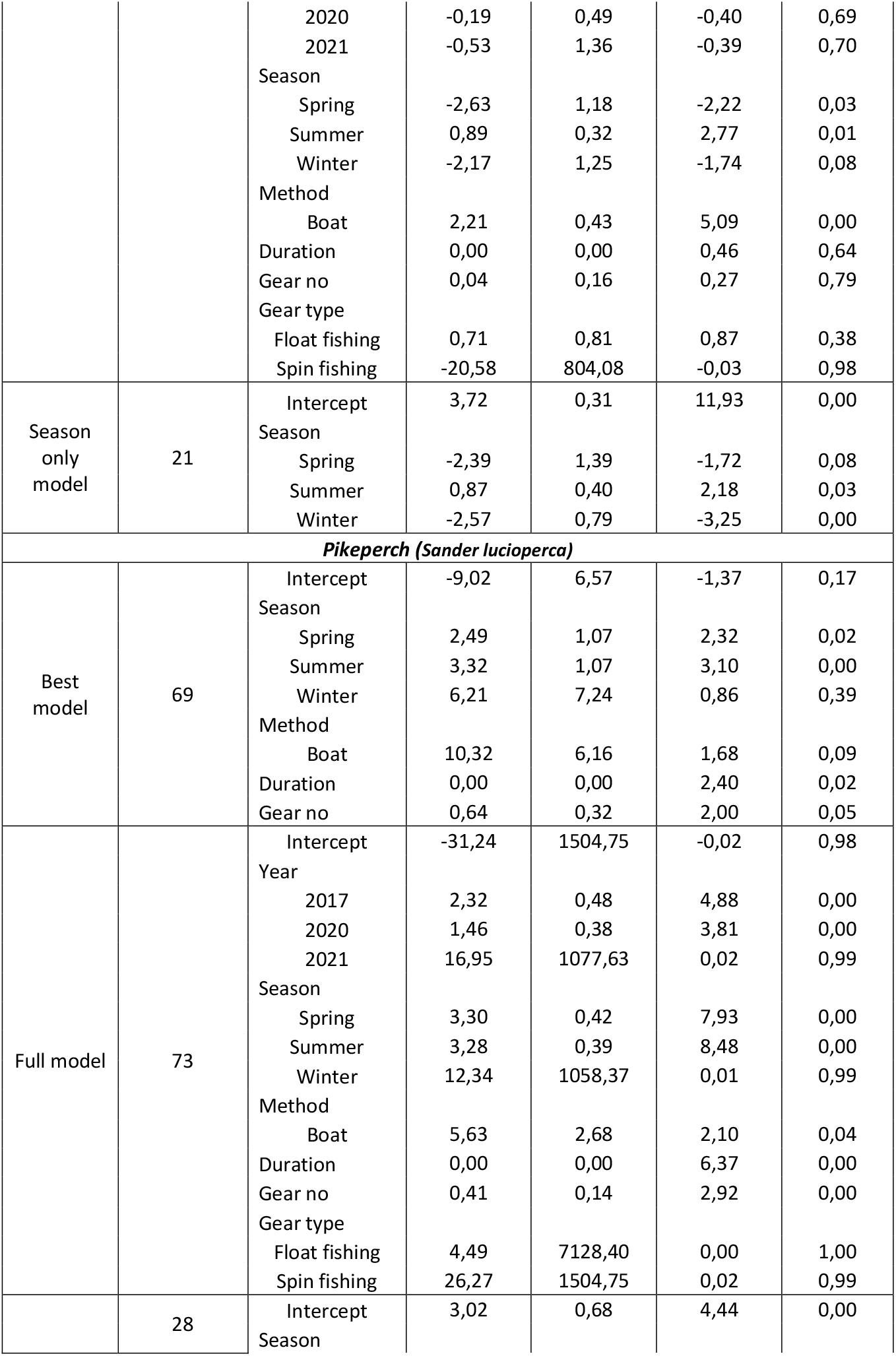
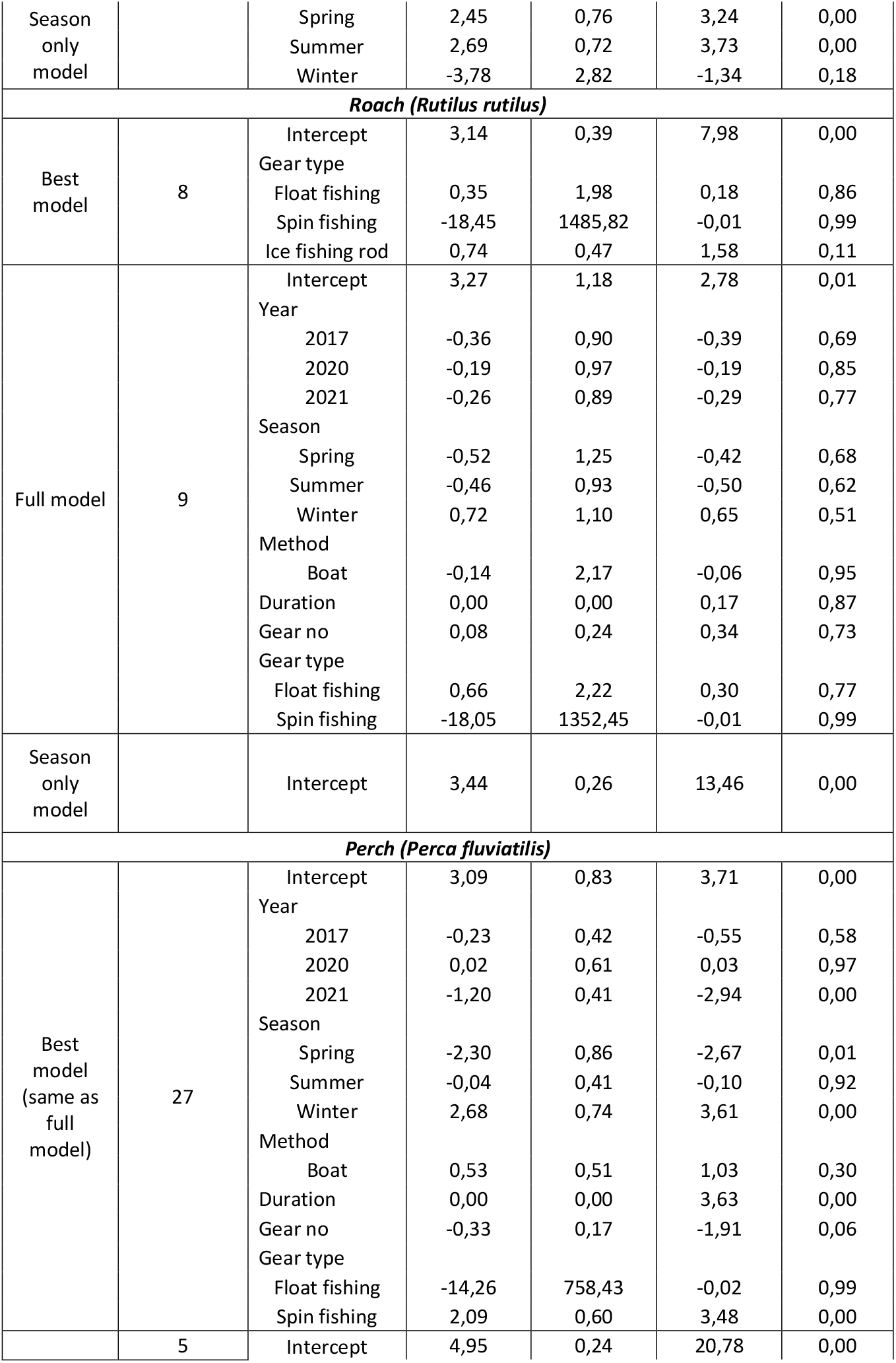
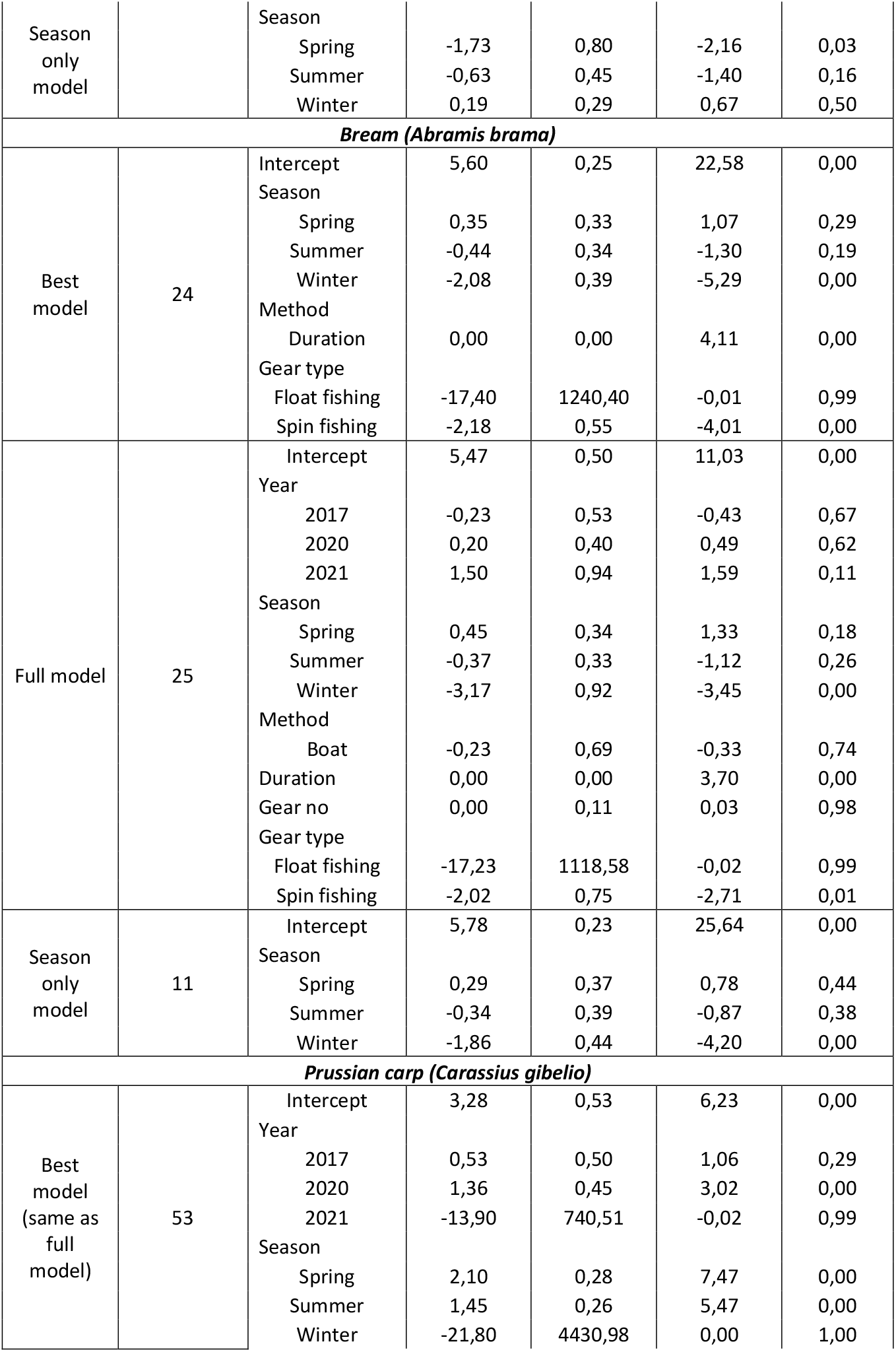
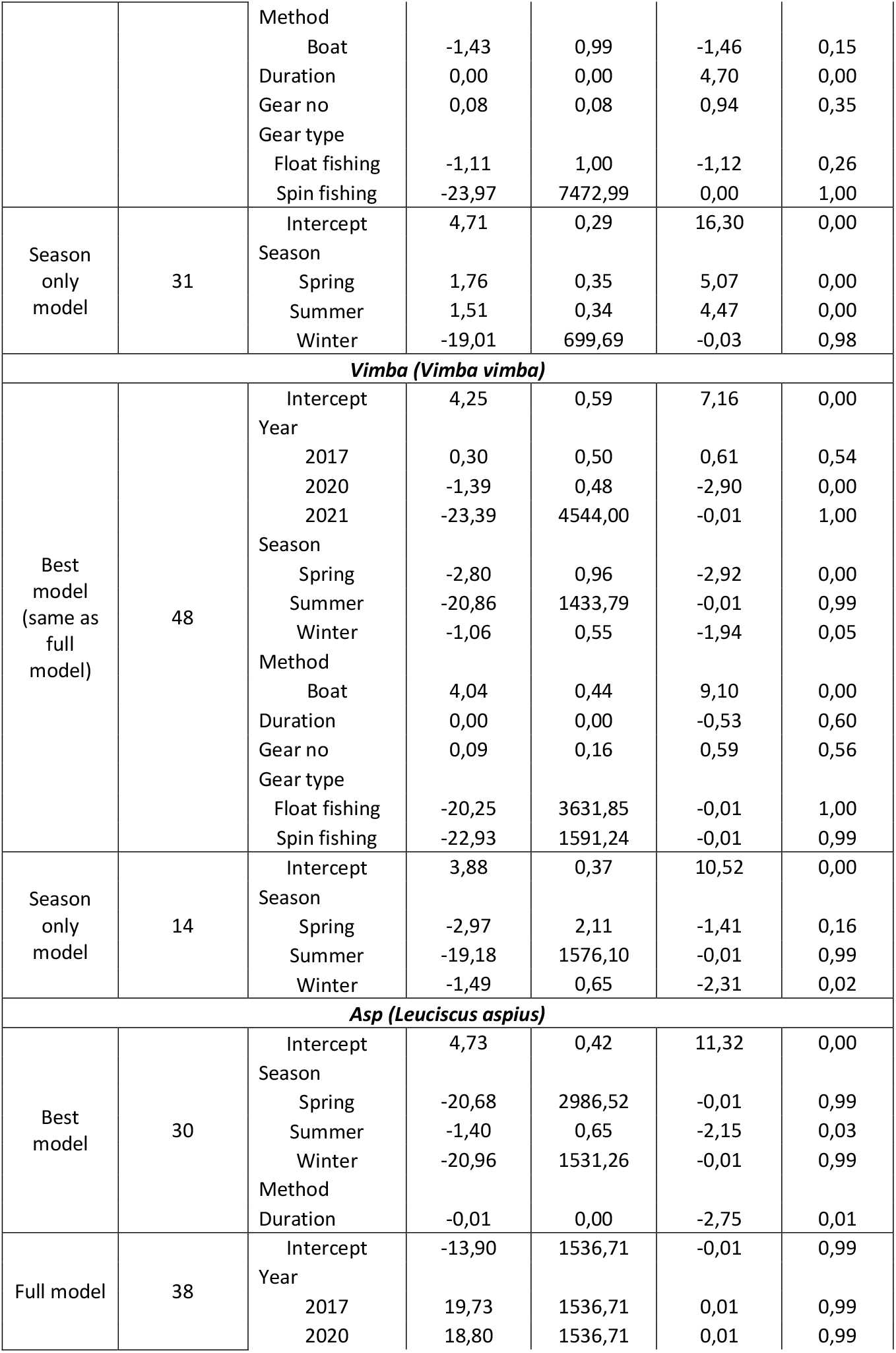
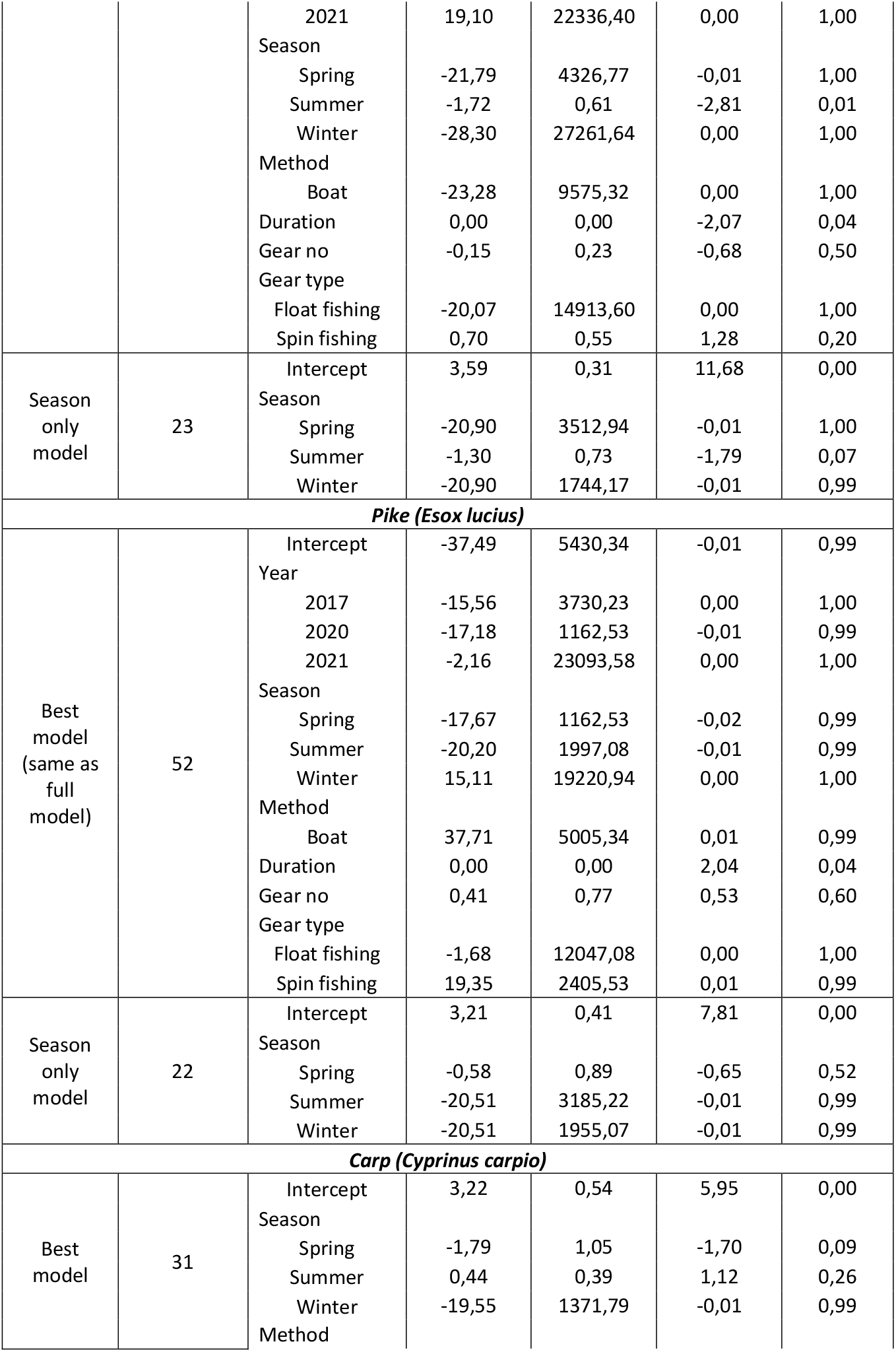
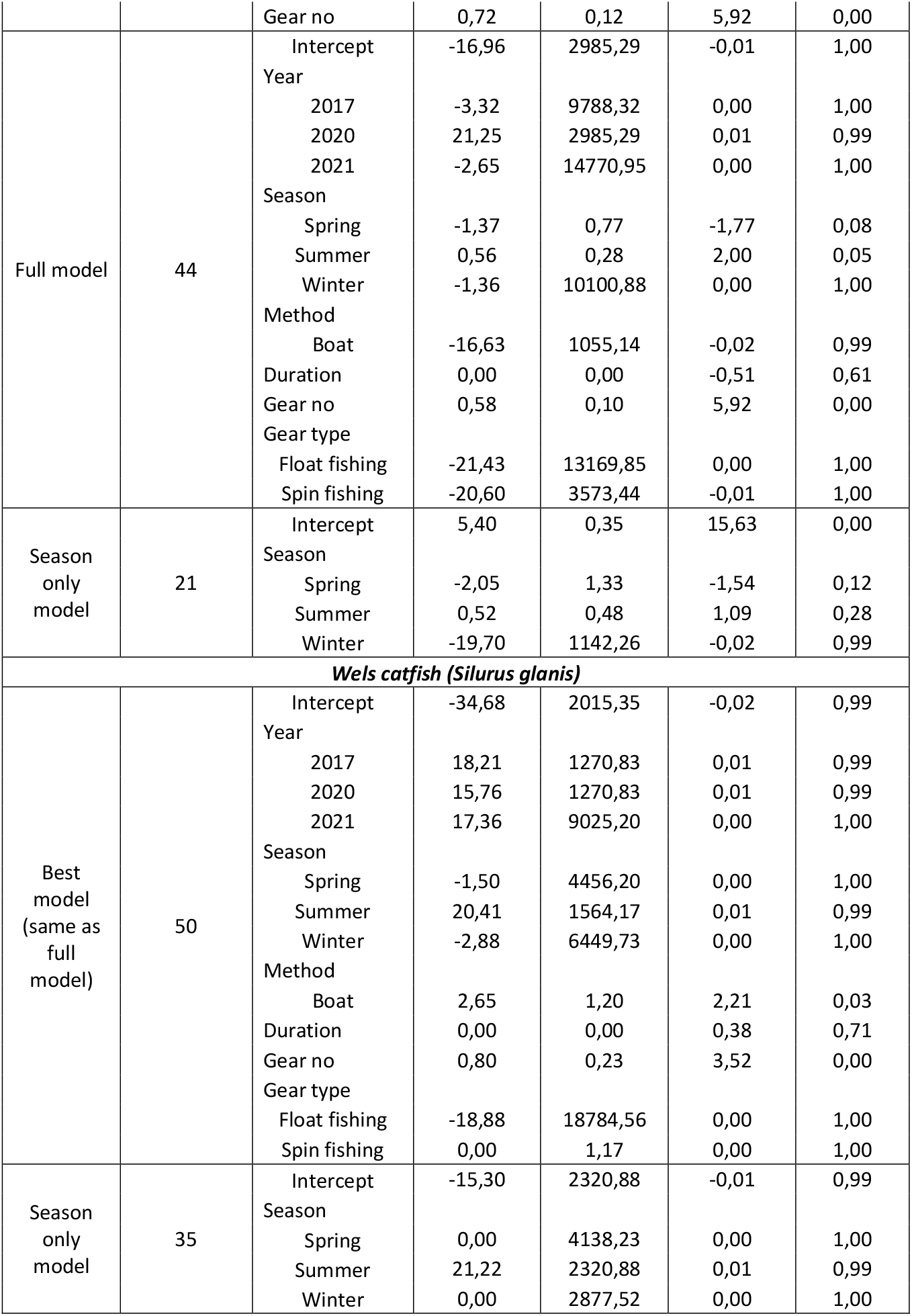
Coefficients of the general linear models applied to recreational fishing catch CPUE estimation. For each species or total catch, we show the full, best and the season only model. Note that for each parameter that is treated as a factor, coefficients are relative to the first value; this is indicated in the coefficients of Total catch. Season “Winter” fully corresponded with the ice fishing season and “Ice fishing rod” gear type. Therefore in most cases only three gear types (Bottom, Spin and Float fishing) are included in the model (except for roach, where season was not included in the best model)

**Supplementary Table A.1.** Parameters included in the best (lowest AIC value) general linear model of scientific CPUE and the proportion of total variance explained. All variables are treated as fixed effects.

|  | Model | Explained deviance (%) |
| --- | --- | --- |
| **All mesh sizes** |  |  |
| **Roach (*Rutilus rutilus*)** | Year + season + Location.gr + Gillnet_category + Net.mesh.gr + Net_length_Or + Soak_time_Or | 42 |
| **Perch (*Perca fluviatilis*)** | Year + season + Location.gr + Gillnet_category + Net.mesh.gr + Net_length_Or + Soak_time_Or | 34 |
| **Pikeperch (*Sander lucioperca*)** | Year + season + Location.gr + Gillnet_category + Net_length_Or + Soak_time_Or | 35 |
| **Bream (*Abramis brama*)** | Year + season + Location.gr + Gillnet_category + Net.mesh.gr + Net_length_Or + Soak_time_Or | 59 |
| **Silver bream (*Blicca bjoerkna*)** | Year + season + Location.gr + Gillnet_category + Net.mesh.gr + Net_length_Or + Soak_time_Or | 29 |
| **Big mesh sizes (>38 mm) only** |  |  |
| **Roach (*Rutilus rutilus*)** | Year + season + Location.gr + Gillnet_category + Soak_time_Or | 38 |
| **Perch (*Perca fluviatilis*)** | Year + season + Location.gr + Gillnet_category + Net_length_Or + Soak_time_Or | 33 |
| **Pikeperch (*Sander lucioperca*)** | Year + season + Location.gr + Gillnet_category + Net_length_Or + Soak_time_Or | 33 |
| **Bream (*Abramis brama*)** | Year + season + Location.gr + Gillnet_category + Net_length_Or + Soak_time_Or | 51 |
| **Silver bream (*Blicca bjoerkna*)** | Year + season + Location.gr + Gillnet_category + Net_length_Or + Soak_time_Or | 25 |

**Supplementary Table A.4.** Estimated recreational catches (grams per fishing trip) and their uncertainty for less common species in Kaunas WR (main species are shown in Table 1).

| Parameter | Mean catch | 2.5% CI | 97.5% CI | *t* | *p* |
| --- | --- | --- | --- | --- | --- |
| **Vimba bream (*Vimba vimba*)** |  |  |  |  |  |
| autumn | 48.44 | 23.48 | 99.93 | 10.52 | 0.00 |
| spring | 2.48 | 0.04 | 145.46 | -1.41 | 0.16 |
| summer | 0.00 | 0.00 | Inf | -0.01 | 0.99 |
| winter | 10.90 | 3.86 | 30.78 | -2.31 | 0.02 |
| **Asp (*Leuciscus aspius*)** |  |  |  |  |  |
| autumn | 36.36 | 19.88 | 66.52 | 11.68 | 0.00 |
| spring | 0.00 | 0.00 | Inf | -0.01 | 1.00 |
| summer | 9.88 | 2.70 | 36.11 | -1.79 | 0.07 |
| winter | 0.00 | 0.00 | Inf | -0.01 | 0.99 |
| **Pike (*Esox lucius*)** |  |  |  |  |  |
| autumn | 24.68 | 11.02 | 55.25 | 7.81 | 0.00 |
| spring | 13.87 | 2.97 | 64.82 | -0.65 | 0.52 |
| summer | 0.00 | 0.00 | Inf | -0.01 | 0.99 |
| winter | 0.00 | 0.00 | Inf | -0.01 | 0.99 |

## Supplementary figures

**Supplementary Figure A.1.**
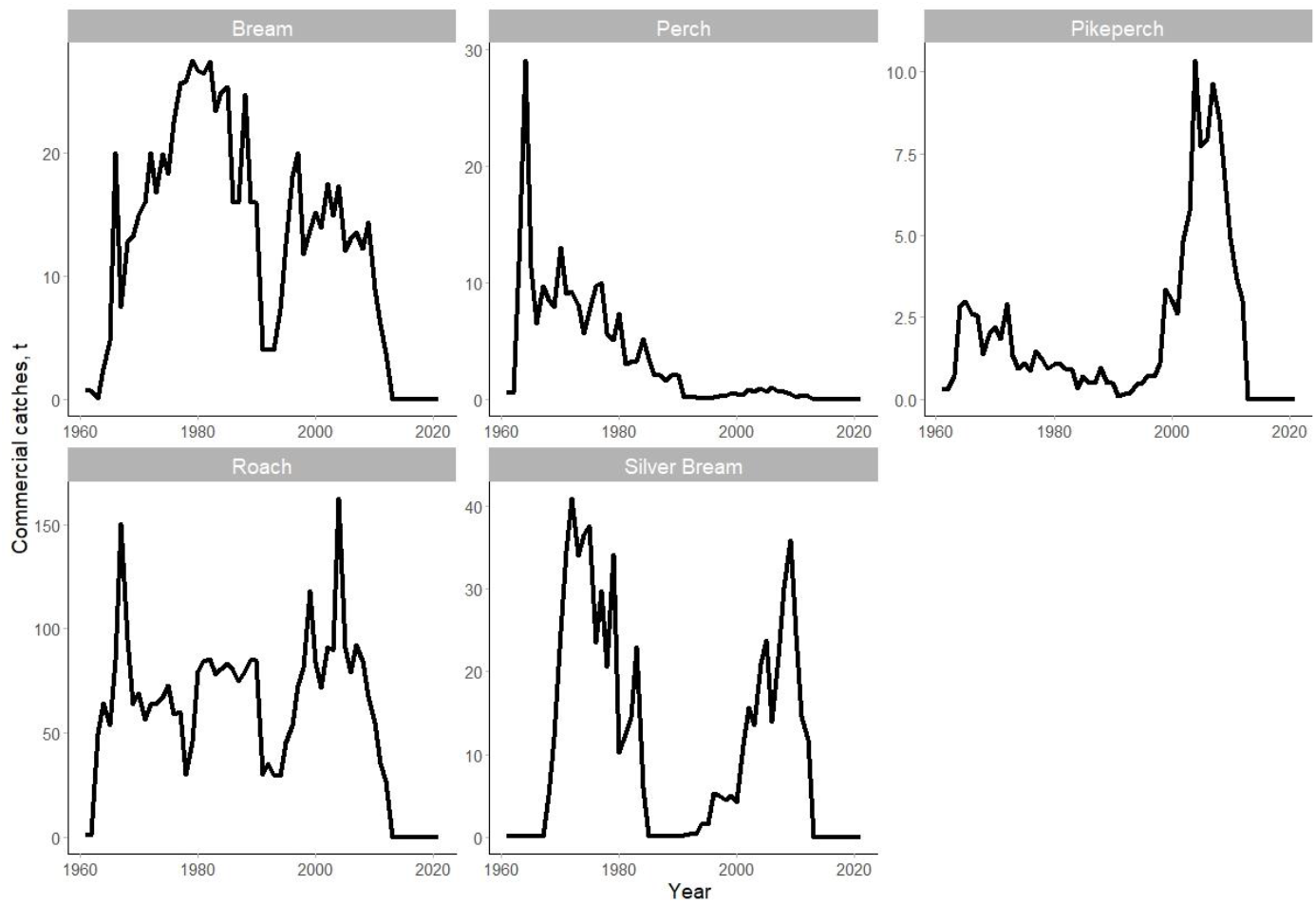
Commercial catches in tonnes (t) since 1961 in Kaunas Reservoir.

**Supplementary Figure A.2.**
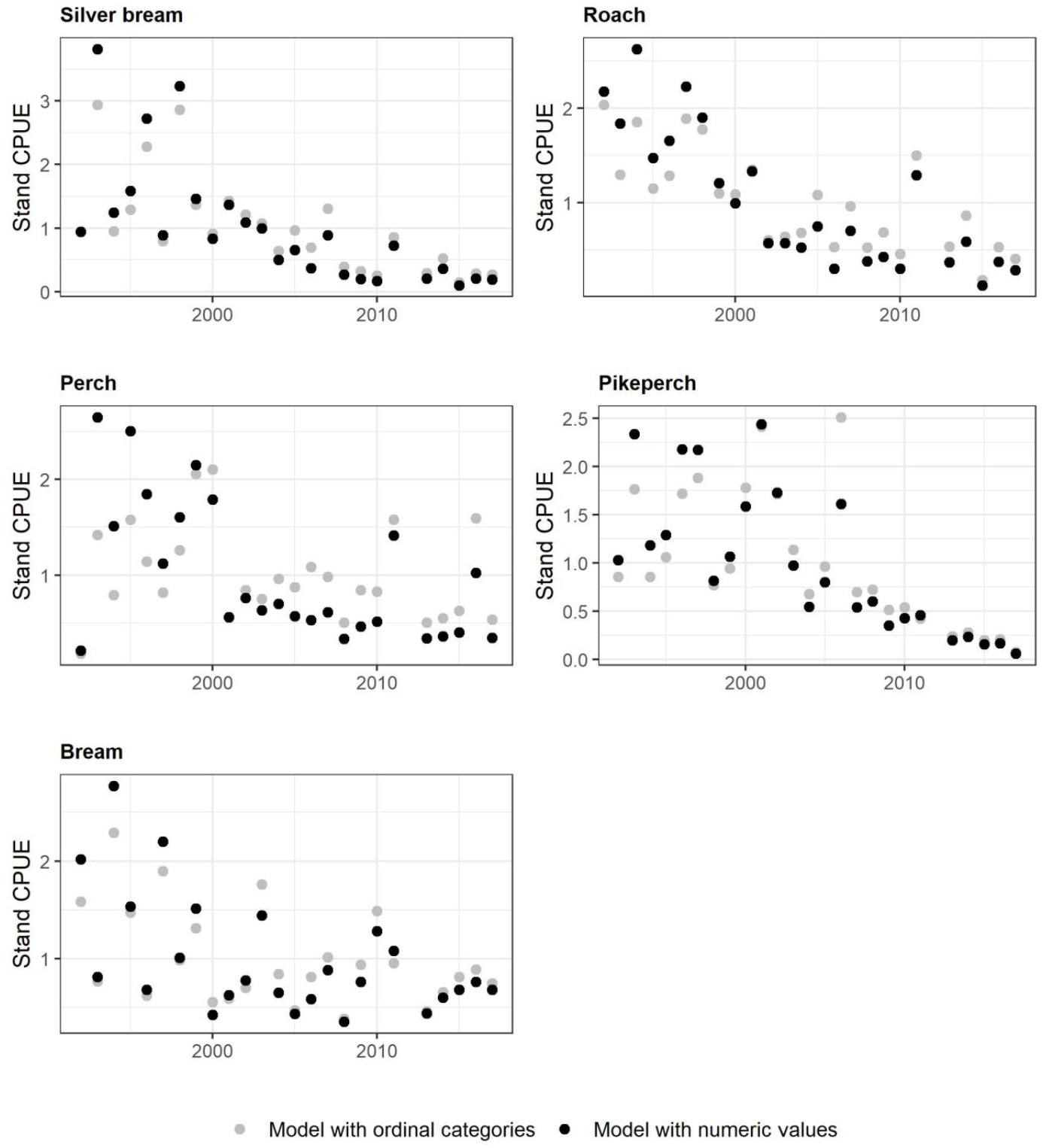
Comparison of the standardized CPUE when treating net length and soak time as numeric vs ordinal variables (Curonian Lagoon data, See Supplementary Table A.2.).

**Supplementary Figure A.3.**
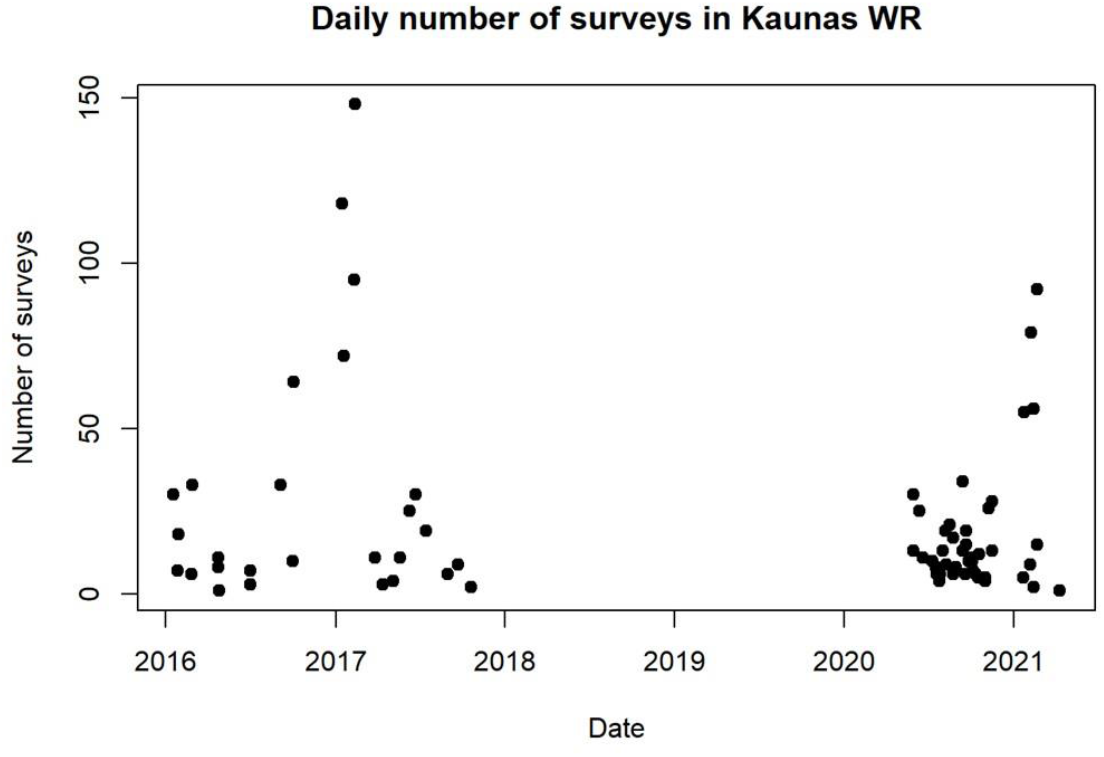
Number of daily angler surveys in Kaunas Reservoir. Days with very high daily survey numbers (> 50) correspond to ice fishing season, when fishing activity is very high.

**Supplementary Figure A.4.**
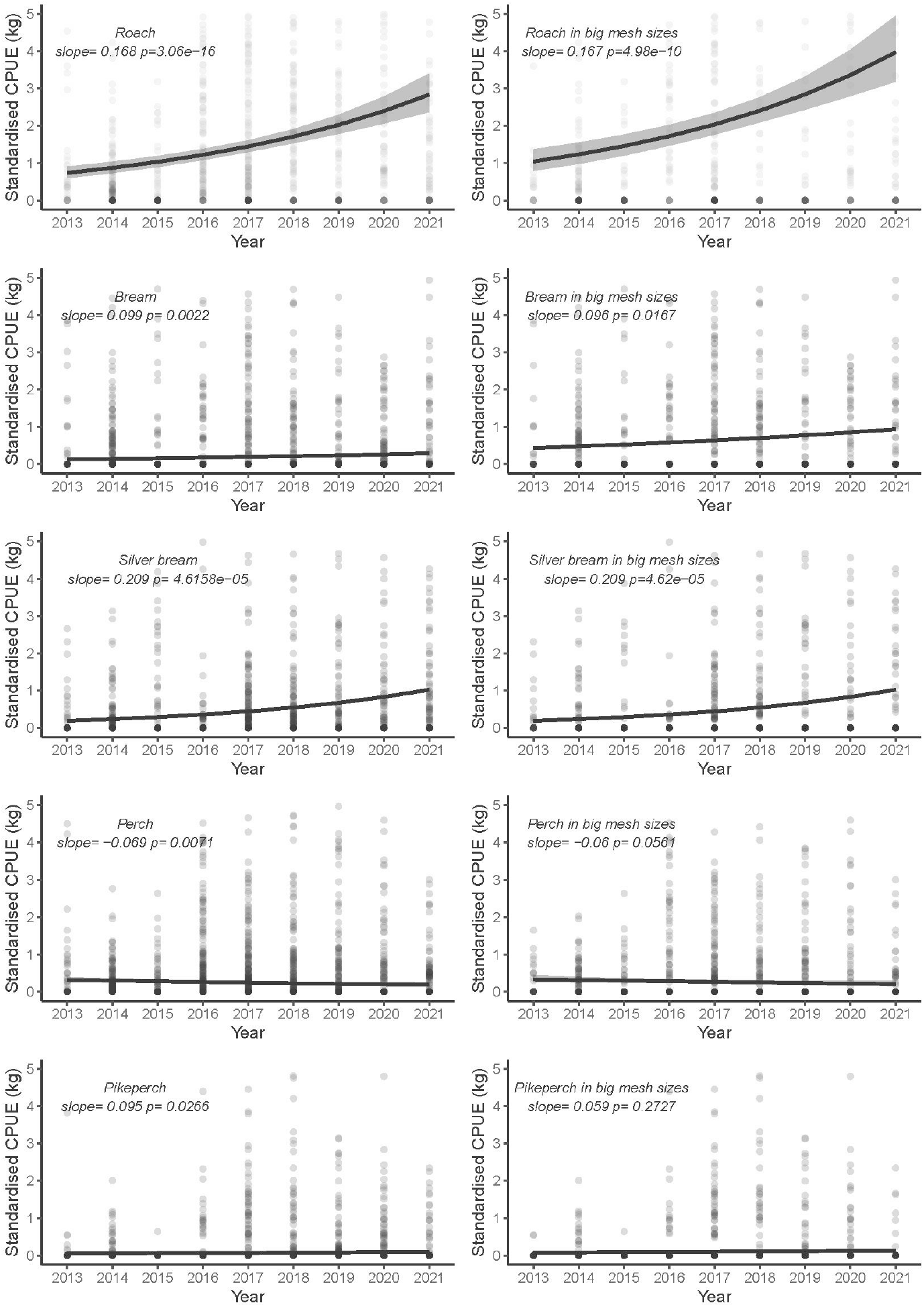
Trends in standardized scientific CPUE since fishery closure across all fish sizes (left) and those restricted to mesh sizes > 38 mm (right).

**Supplementary Figure A.5.**
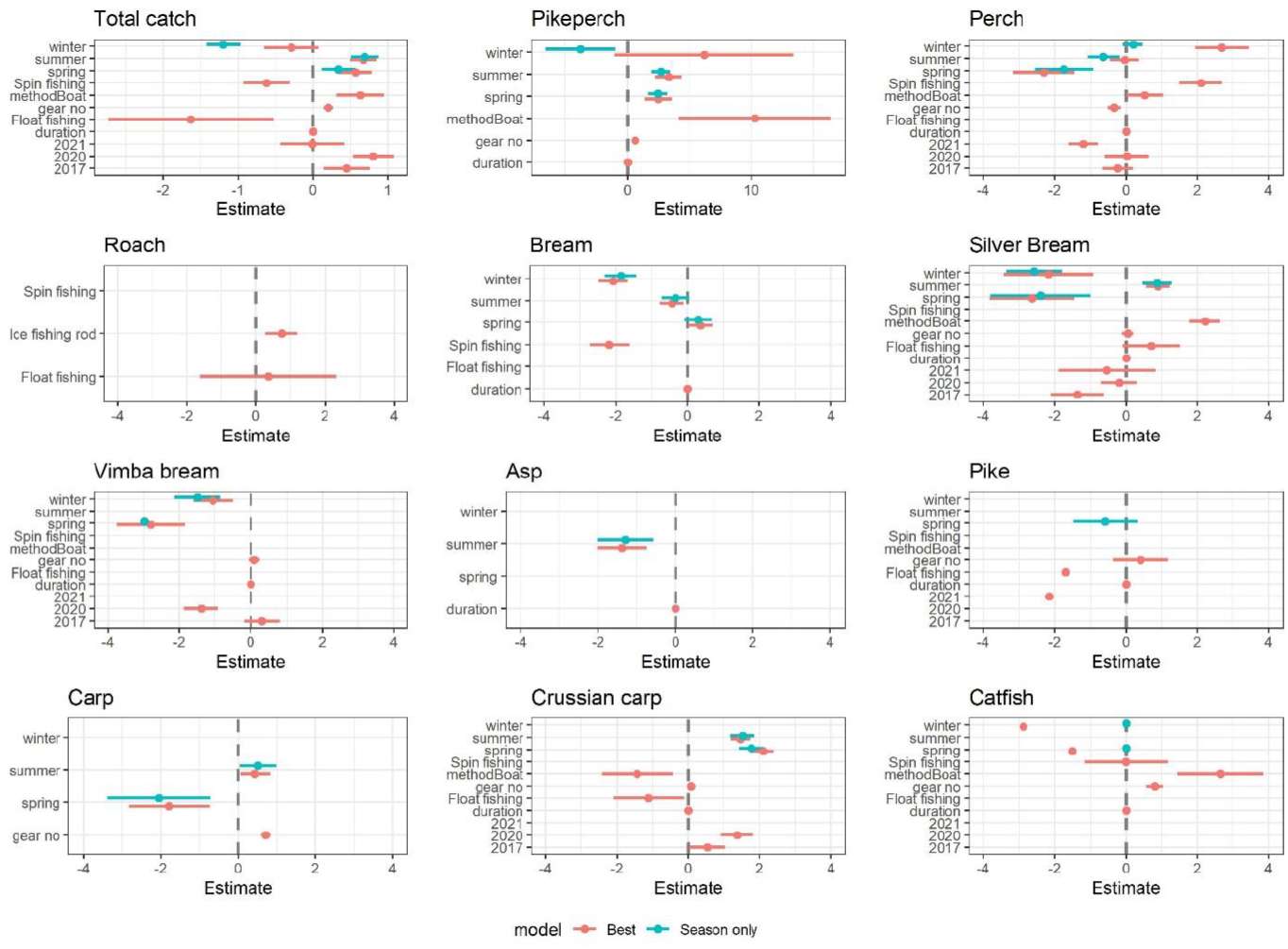
Estimates of general linear model coefficients (± standard error) for total and species-specific recreational catch. Coefficient estimates for the best model are shown in red and those for the season-only model (used to predict total angler catches) are shown in blue. Due to high confidence intervals, values of some estimates are not shown as they are out of scale (provided in Table Sx). The coefficients for variables that are treated as factors in the GLM are shown as differences from the first value of each parameter; they are *autumn* for ‘season’, *Feeder* for ‘gear_type’ (compared to Spin and Float fishing), *shore fishing* for ‘method’, and *2016* for ‘year’; ‘gear_no’ and ‘fishing_duration’ were treated as numeric variables; and ‘fishing_duration’ was estimated in minutes. The coefficient values were very small, despite being highly significant in many cases (see Table A.4.).

## References

1. Abell R., Allan J.D., Lehner B. 2007. Unlocking the potential of protected areas for freshwaters. Biological Conservation, 134(1):48–63, https://doi.org/10.1016/j.biocon.2006.08.017.

2. Ahrens R.N., Allen M.S., Walters C., & Arlinghaus R. 2020. Saving large fish through harvest slots outperforms the classical minimum-length limit when the aim is to achieve multiple harvest and catch-related fisheries objectives. Fish and Fisheries, 21(3), 483–510.

3. Arlinghaus R., Tillner R., Bork M. 2015. Explaining participation rates in recreational fishing across industrialised countries. Fisheries Management and Ecology, 22(1), 45–55. doi:10.1111/fme.12075

4. Arlinghaus R. and Cooke S.J. 2009. Recreational Fisheries: Socioeconomic Importance, Conservation Issues and Management Challenges. In Recreational Hunting, Conservation and Rural Livelihoods (eds B. Dickson, J. Hutton and W.M. Adams). https://doi.org/10.1002/9781444303179.ch3

5. Barnes M. 2014. A Comparison of A Creel Census to Modeled Access-Point Creel Surveys on Two Small Lakes Managed as Put-and-Take Rainbow Trout Fisheries. Fisheries and Aquaculture Journal. 05. 10.4172/2150-3508.1000086.

6. Bartholomew & Bohnsack J.A. 2005. A review of catch-and-release angling mortality with implications for no-take reserves. Reviews in Fish Biology and Fisheries (2005) 15: 129–154. DOI 10.1007/s11160-005-2175-1.

7. Beard T.D. Jr., Essington T.E. 2000. Effects of angling and life history processes on bluegill size structure: insights from an individual-based model. Transactions of the American Fisheries Society 129: 561–568.

8. Beard T.D. Jr., Kampa J.M. 1999. Changes in bluegill, black crappie, and yellow perch populations in Wisconsin during 1967–1991. North American Journal of Fisheries Management 19:1037–1043.

9. Bennett B.A. 1991. Long-term trends in the catches by shore anglers in False Bay. Trans. R. Soc. S. Afr. 47, 683–692.

10. Coleman F.C, Figueira W.F., Ueland J.S., Crowder L.B. 2004. The Impact of United States Recreational Fisheries on Marine Fish Populations. Science 305(5692):1958–60.

11. Conron S., Bridge N.F., Oliveiro P., Bruce T.K. 2012. Angler diary monitoring of recreational fishing in selected Victorian waters during 2010/11. Recreational Fishing Grant Program Final Report. Department of Primary Industries, Victoria, Australia. 24 pp.

12. Cooke S.J., Cowx I.G. 2004. The Role of Recreational Fishing in Global Fish Crises. Bioscience 54(9): 857–59.

13. Cooke S.J., Cowx I.G. 2006. Contrasting Recreational and Commercial Fishing: Searching for Common Issues to Promote Unified Conservation of Fisheries Resources and Aquatic Environments. Biological Conservation 128(1):93–108.

14. Cooke S., Schreer J., Wahl, D., Philipp D. 2002. Physiological impacts of catch-and-release angling practices on largemouth and smallmouth bass. American Fisheries Society Symposium. 31.

15. Cowx I.G. 2002. Analysis of Threats to Freshwater Fish Conservation: Past and Present Challenges. In: Collares-Pereira, M.J., Cowx, I.G. and Coelho, M.M., Eds., Conservation of Freshwater Fish: Options for the Future, Blackwell Science, Oxford, 201–220.

16. Cowx I.G. 2015. Characterisation of inland fisheries in Europe. Fish Manag Ecol, 22: 78–87. https://doi.org/10.1111/fme.12105

17. Dainys J., Mateos-González F., Skov C., Urbonavičius R., Audzijonyte A. Angling counts: harnessing the power of technological advances for recreational fishing surveys. (Under review, see supplement).

18. Dunn P.K. 2017. Tweedie: Evaluation of Tweedie exponential family models. R package version 2.3.3.

19. FAO. 1999. Review of the State of World Fishery Resources: Inland Fisheries. Rome: FAO Fisheries Department. FAO Fisheries Circular no. 942.

20. FAO. 2010. The State of World Fisheries and Aquaculture 2010. FAO Fisheries and Aquaculture Department, Rome. 214 p.

21. FAO. 2017. The role of Recreational Fisheries in the sustainable management of marine resources pp. 65 – 68 In, GLOBEFISH Highlights - Issue 2/2017, FAO Rome 80 pp.

22. Forrestal F., Goodyear C., Schirripa M. 2019. Applications of the longline simulator (LLSIM) using US pelagic longline logbook data and Atlantic blue marlin. Fisheries Research. 211. 331–337. 10.1016/j.fishres.2018.11.029.

23. Funge-Smith S., Bennett A. 2019. A fresh look at inland fisheries and their role in food security and livelihoods. Fish Fish. 20(6): 1176–1195. https://doi.org/10.1111/faf.12403

24. Giner G., Smyth, G.K. 2016. statmod: probability calculations for the inverse Gaussian distribution. R Journal 8(1), 339–351.

25. Haddon. 2020. r4cpue: Functions to assist with CPUE standardization. R package version 0.1.3.

26. Hilborn R., Branch T., Ernst B., Magnusson A., Minte-Vera C., Scheuerell M., Valero J. 2003. State of the World’s Fisheries. Annual Review of Environment and Resources. 28. 359–399. 10.1146/annurev.energy.28.050302.105509.

27. Jones C.M., Robson D.S., Lakkis H.D., Kressel J. 1995. Properties of catch rates in analysis of angler surveys. Transactions of the American Fisheries Society 124: 911–928.

28. Kaemingk M., Chizinski C.J., Hurley K.L., Pope K.L. 2018. Synchrony—An emergent property of recreational fisheries. Journal of Applied Ecology 55:2986–2996.

29. Kearney R.E. Fisheries property rights and recreational/commercial conflict: Implications of policy development in Australia and New Zealand. Marine Policy 2001; 25:49–59.

30. Kristensen E., Sand-Jensen K., Martinsen K.T., Madsen-Østerbye M., Kragh T. 2020. Fingerprinting pike: The use of image recognition to identify individual pikes. Fisheries Research. https://doi.org/10.1016/j.fishres.2020.105622.

31. Kummu M., de Moel H., Ward P.J., Varis O. 2011. How close do we live to water? A global analysis of population distance to freshwater bodies. PLoS One, 6(6): e20578. doi:10.1371/journal.pone.0020578. PMID:21687675.

32. Kura Y., Revenga C., Hoshino E., Mock G. 2004. Fishing for Answers: Making Sense of the Global Fish Crisis.

33. Lewin W.C., Arlinghaus R., Mehner T. 2006. Documented and Potential Biological Impacts of Recreational Fishing: Insights for Management and Conservation, Reviews in Fisheries Science, 14(4): 305–367, https://doi.org/10641260600886455

34. Lockwood R.N., Benjamin D.M., Bence J.R. 1999. Estimating angling effort and catch from Michigan roving and access site angler survey data. Michigan Department of Natural Resources, Fisheries Research Report 2044.

35. Ložys L., Stanevičius V., Pūtys Ž., Dainys J., Levickienė D., Jakimavičius D., Akstinas V., Adžgauskas G., Tomkevičienė A., Irbinskas V. 2020. Assessment of the impact of water level fluctuations on fish and waterbird populations in Kaunas Reservoir. 98p.

36. Mallison C.T., Cichra C.E. 2004. Accuracy of angler-reported harvest in roving creel surveys. North American Journal of Fisheries Management 24: 880–889.

37. Malvestuto S.P. 1996. Sampling the recreational creel. (2^nd^ edn), Pages 591–623 in B. R. Murphy and D. W. Willis, editors. Fisheries Techniques, American Fisheries Society, Bethesda, Maryland.

38. Mangin T., Costello C., Anderson J., Arnason R., Elliott M., et al. 2018. Are fishery management upgrades worth the cost?. PLOS ONE 13(9):e0204258. https://doi.org/10.1371/journal.pone.0204258

39. Manning R.E. 2010. Studies in outdoor recreation: Search and research for satisfaction. Oregon State University Press.

40. McPhee D., Leadbitter D., Skilleter A.G. 2002. Swallowing the bait: is recreational fishing in Australia ecologically sustainable? Pacific Conservation Biology 8(1), 40–51. doi: https://doi.org/10.1071/PC020040.

41. Morales-Nin B., Moranta J., García C., Tugores M.P., Grau A.M., Riera F., Cerdà M. 2005. The recreational fishery off Majorca Island (western Mediterranean): some implications for coastal resource management, ICES Journal of Marine Science, 62(4): 727–739, https://doi.org/10.1016/j.icesjms.2005.01.022

42. Muoneke M.I., W. Childress M. 1994. Hooking mortality: A review for recreational fisheries, Reviews in Fisheries Science, 2(2): 123–156. DOI: 10.1080/10641269409388555

43. Olson D.E., Cunningham P.K., 1989. Sport-fishing trends shown by an annual Minnesota fishing contest over a 58-year period. N. Am. J. Fish. Manage. 9: 287–297.

44. Papenfuss J.T., Phelps N., Fulton, D., Venturelli P.A. 2015. Smartphones Reveal Angler Behavior: A Case Study of a Popular Mobile Fishing Application in Alberta, Canada. Fisheries 40(7): 318–327. 10.1080/03632415.2015.1049693,

45. Persson L. 1987. Effects of habitat and season on competitive interactions between roach *(Rutilus rutilus)* and perch *(Perca fluviatilis)*. Oecologia (Berlin) 73:170–177

46. Pope K.L., Powel L.A., Harmon B.S., Pegg M.A., Chizinski C.J. 2017. Estimating the number of recreational anglers for a given waterbody a given waterbody. Fisheries Research 191: 69–75 DOI: http://dx.doi.org/10.1016/j.fishres.2017.03.004

47. Post J.R., Sullivan M., Cox S., Lester N.P., Walters C.J., Parkinson E.A., Paul A.J., Jackson L., Shuter B.J. 2002. Canada’s recreational fisheries: the invisible collapse? Fisheries 27: 6–17.

48. Shono H. 2008. Confidence interval estimation of CPUE year trend in delta-type two-step model. Fisheries Science 74: 712–717.

49. Stoll-Kleemann S., de la Vega-Leinert A., Schultz L. 2010. The role of community participation in the effectiveness of UNESCO Biosphere Reserve management: Evidence and reflections from two parallel global surveys. Environmental Conservation. 37: 227–238. 10.1017/S037689291000038X.

50. Su M., Wang L., Xiang J., et al. 2021. Adjustment trend of China’s marine fishery policy since 2011. Marine Policy. 124: 104322.

51. Vandergoot C.S. 2014. Estimation of regional mortality rates for Lake Erie walleye *Sander vitreus* using spatial tag-recovery modeling. PhD thesis, Michigan State University.

52. VanDeValk A.J., Jackson J.R., Krueger S.D., Brooking T.E., Rudstam L.G. 2007. Influence of party size and trip length on angler catch rates on Oneida Lake, New York. North American Journal of Fisheries Management 27: 127–136.

53. Vejřík L., Matějíčková I., Sed’a J., Blabolil P., Jůza T., et al. 2016 Who Is Who: An Anomalous Predator-Prey Role Exchange between Cyprinids and Perch. PLOS ONE 11(6): e0156430. https://doi.org/10.1371/journal.pone.0156430

54. Venohr M., Langhans S.D., Peters O., Hölker F. Arlinghaus R., Mitchell L., Wolter C. 2018. The underestimated dynamics and impacts of water-based recreational activities on freshwater ecosystems. Environmental Reviews 26(2): 199–213. https://doi.org/10.1139/er-2017-0024

55. Wadiwel D. 2019. ‘Fishing for Fun’: The Politics of Recreational Fishing, Animal Studies Journal, 8: 202–228.

56. Walters C. 1998. Designing fisheries management systems that do not depend upon accurate stock assessment. In Reinventing fisheries management (pp. 279–288). Springer, Dordrecht.

57. West R.J., Gordon G.N.G. 1994. Commercial and recreational harvest of fish from two Australian coastal rivers. Australian Journal of Marine and Freshwater Research 45, 1259–1279.

58. Winstanley R. 2019. A wake-up call for recreational fishing in Australia. SETFIA News 22nd February 2019 Accessed at https://setfia.org.au/a-wake-up-call-for-recreational-fishing-in-australia/ on 17-Apr-2021.

59. Yount J. D. 1991. Ecology and management of the Zebra Mussel and other introduced aquatic nuisance species.

60. Zeller D., Darcy M., Booth S., Lowe M.K., Martell S. 2008. What about recreational catch?: Potential impact on stock assessment for Hawaii’s bottomfish fisheries? Fisheries Research, 91(1): 88–97. https://doi.org/10.1016/j.fishres.2007.11.010

61. Zhang H., Wu J., Gorfine H., Shan X., Shen L., Yang H., Du H., Li J., Wang C., Zhou Q., Liu Z., Kang M., Wei Q. 2020. Inland fisheries development versus aquatic biodiversity conservation in China and its global implications. Reviews in Fish Biology and Fisheries. 30: 637–655. https://doi.org/10.1007/s11160-020-09622-y

